# Survey on Open Science Practices in Functional Neuroimaging

**DOI:** 10.1101/2021.11.26.470115

**Authors:** Christian Paret, Nike Unverhau, Franklin Feingold, Russell A. Poldrack, Madita Stirner, Christian Schmahl, Maurizio Sicorello

**Affiliations:** Department of Psychosomatic Medicine and Psychotherapy, Central Institute of Mental Health Mannheim, Medical Faculty Mannheim / Heidelberg University, Germany.; Sagol Brain Institute, Wohl Institute for Advanced Imaging, Tel-Aviv Sourasky Medical Center and School of Psychological Sciences, Tel-Aviv University, Israel; Department of Psychology, Stanford University, Stanford, CA

**Author notes:** Correspondence to: Christian Paret, Central Institute of Mental Health J5, D-68159 Mannheim, Germany.

**Keywords:** data sharing, fMRI, metascience, neuroimaging, open science, preregistration, research methods, replication, reproducibility, robustness, validity

## Abstract

Replicability and reproducibility of scientific findings is paramount for sustainable progress in neuroscience. Preregistration of the hypotheses and methods of an empirical study before analysis, the sharing of primary research data, and compliance with data standards such as the Brain Imaging Data Structure (BIDS), are considered effective practices to secure progress and to substantiate quality of research. We investigated the current level of adoption of open science practices in neuroimaging and the difficulties that prevent researchers from using them.

Email invitations to participate in the survey were sent to addresses received through a PubMed search of human functional magnetic resonance imaging studies between 2010 and 2020. 283 persons completed the questionnaire.

Although half of the participants were experienced with preregistration, the willingness to preregister studies in the future was modest. The majority of participants had experience with the sharing of primary neuroimaging data. Most of the participants were interested in implementing a standardized data structure such as BIDS in their labs. Based on demographic variables, we compared participants on seven subscales, which had been generated through factor analysis. It was found that experienced researchers at lower career level had higher fear of being transparent, researchers with residence in the EU had a higher need for data governance, and researchers at medical faculties as compared to other university faculties reported a higher need for data governance and a more unsupportive environment.

The results suggest growing adoption of open science practices but also highlight a number of important impediments.

## 1 Introduction

Neuroimaging, and in particular functional magnetic resonance imaging (fMRI), has contributed greatly to the generation and testing of neural models of brain function and dysfunction in mental disorders. Although the number of neuroimaging publications increases with every year, a growing literature is shaking the ground, questioning the replicability of many reported findings ^1–4^. Assessing validity requires researchers to be fully transparent about the *a priori* hypotheses underlying a study, the complete reporting of methods, and the availability of data to reproduce the findings. These conditions are often not met ^5, 6^. Open science practices can protect against such adversities, but they confront scientists with additional demands to learn and adopt new techniques. To accelerate the implementation of open science practices, it is necessary to better understand obstacles that prevent researchers from adopting these practices. While survey data are available on researchers’ preferences, barriers and fears related to data sharing in psychology ^7^, open science practices besides data sharing have not been surveyed in the behavioral sciences, yet. Neuroimaging data is complex and hard to de-identify ^8–12^, confronting researchers in this field with intricate challenges to share data. We investigated the familiarity, adoption, experience, and obstacles concerning open science practices in neuroimaging research. We focused on three fundamental instruments of a reproducible science: Preregistration, data sharing, and current standards of formatting and structuring data as implemented with the Brain Imaging Data Structure (BIDS)^13^. In a preregistration, authors provide an overview on the planned study and explain the *a priori* hypotheses along with the methods they plan to use to test the hypotheses ^14^. The document is time-stamped and any changes made thereafter are documented for transparency. Preregistrations are instrumental to avoid confusion of *a priori* and a *posteriori* definition of hypotheses and analysis methods, which can easily lead to flawed interpretation of a p-value from a statistical result and can create overconfidence in findings ^15–17^. In face of high flexibility in preprocessing and analysis methods ^1, 18^, preregistration can dramatically enhance the transparency of a neuroimaging project. More than in basic science, it is mandatory to register clinical trials in a public registry before data acquisition, in order to publish in a renowned biomedical journal. In practice, leading clinical registries leave it at the discretion of the researcher as to how much detail they use to describe the analytic strategy for processing their neuroimaging data. One may register a neuroimaging endpoint in some way similar to “higher BOLD response in ROI (Region-of-Interest) X for the contrast of conditions A vs. B”. There are many possible analysis strategies to assess this endpoint; the search space for significant voxels could be extended to the whole-brain or reduced to a small volume defined by a ROI mask, the mask could be anatomically or functionally defined, and so on. For a confirmatory hypothesis test, the complete analysis plan should be defined *a priori* ^2, 19^, but this is hardly the case in clinical trials with neuroimaging endpoints.

A growing literature is providing tools and guidelines to facilitate reproducible neuroimaging findings and data sharing ^2, 20–23^. Standards such as BIDS, which was introduced by Gorgolewski et al. in 2016 ^13^, present a well-documented scheme to structure data files in directories, provide agreed upon terminology for naming these files, and explain how metadata should be reported. The sharing of primary research data is critical for a reproducible science and can save resources, as existing data can be re-used and aggregated with other data sets for future research projects. Still, researchers often eschew data sharing, e.g. because of a lack of incentives, the fear of misuse, and legal issues such as data protection and privacy issues ^7, 24–27^. In this respect, it is of interest how the General Data Protection Regulation (GDPR), which came into force in the European Union (EU) May, 2018, may affect the preference to share data among researchers inside vs. outside the EU. Moreover, the more complex the dataset, the more resources may be required to prepare a sharable dataset, thus taking up time that could be used to do new experiments ^28^. Where scientist practitioners must balance research and clinical work and where data are collected from vulnerable patient populations, the situation can be even more fraught. Therefore, we analyzed differences between researchers who indicated an affiliation with a medical faculty vs. a different, non-medical faculty. Data and materials from this research are available online ^29^.

## 2 Methods

### 2.1 Participants

A PubMed search with the search terms (“fMRI” OR “functional magnetic resonance imaging” OR “functional Magnetic Resonance Imaging”) was done to collect email addresses from corresponding authors of scientific articles published between 2010/01/01 and 2020/08/28. The “Humans” search filter was applied to exclude animal imaging work. An email was sent to 14,690 addresses on 2020/01/12 with an invitation to participate, including a personalized link to the survey. If the recipients did not click the link or did not complete the survey after 14 days, they received a single reminder email. Figure 1 illustrates the recruitment approach. From 342 persons who clicked the invitation link, 82.75% completed the questionnaire and were included in analysis, corresponding to an overall response rate of 2.42 % and resulting in N=283 participants to-be-analyzed. It took participants 9.62 min on average (3.17; numbers reported in brackets are standard deviations) to arrive at the final slide. Participants were aged 43.89 years on average (9.74), dominantly male (66 %), mostly trained in psychology (40%) (Figure 2), and reported an average research experience of 16.58 years (8.49). Most were affiliated with a university (Figure 3) and reported themselves in cognitive neuroscience (Figure 4). Participants from the European Union were overrepresented in the sample, while the USA and UK ranked second and third in number of participants (Figure 5). Half of the sample held a full or associate professorship or a comparable position (Figure 6).

**Figure 1.**
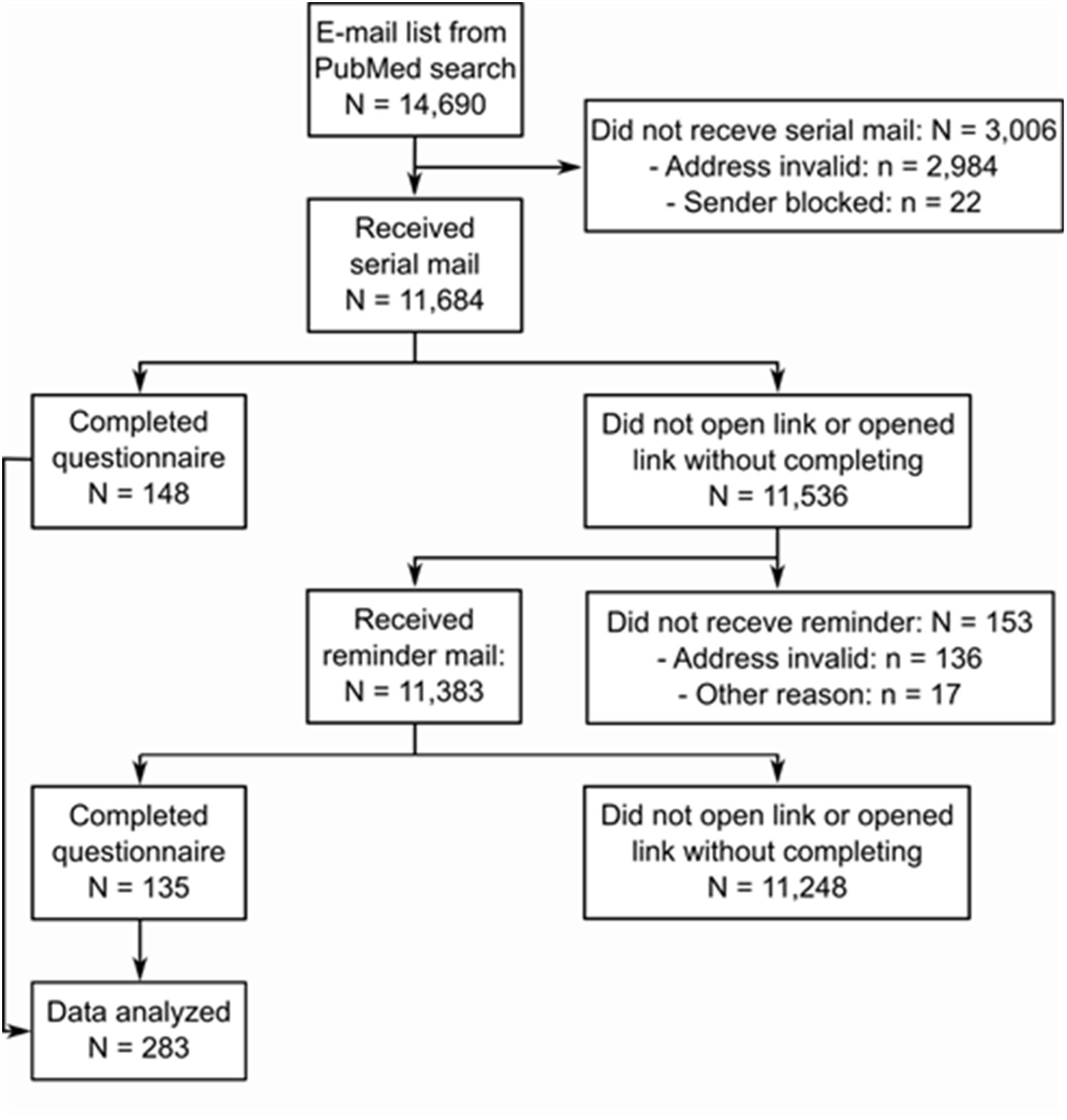
Summary of recruitment approach and number of responses at each step

**Figure 2.**
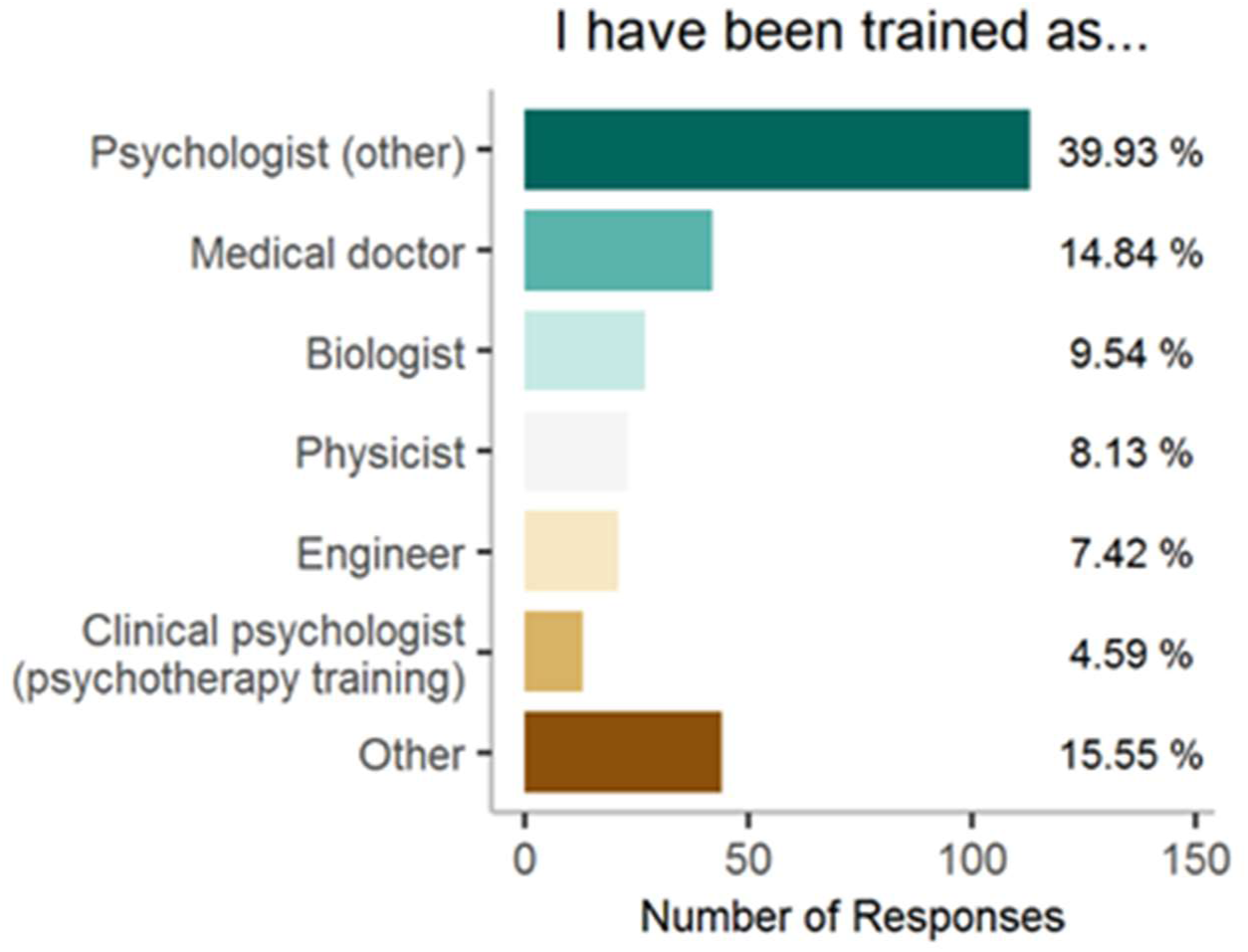
Professional training

**Figure 3.**
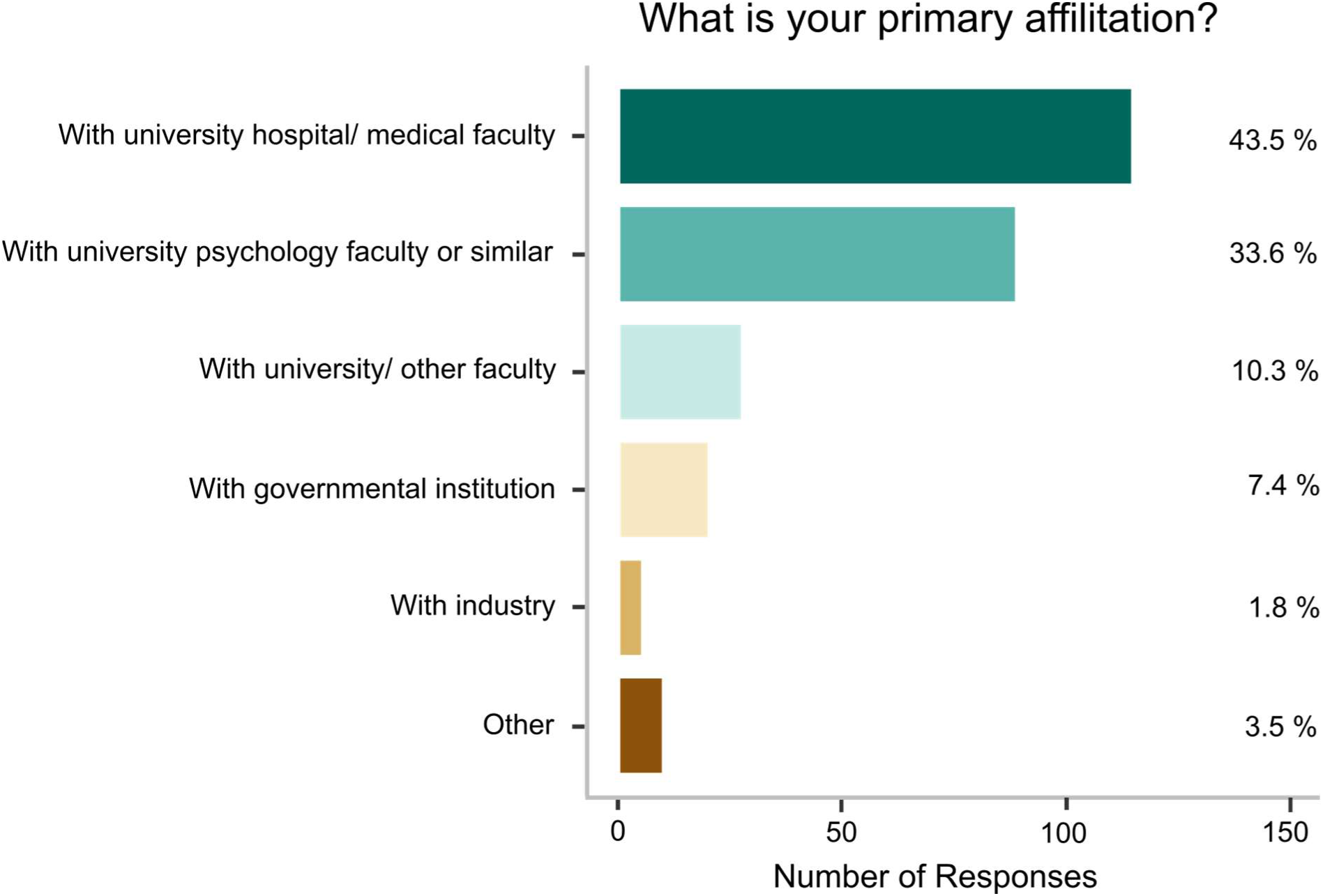
Primary affiliation

**Figure 4.**
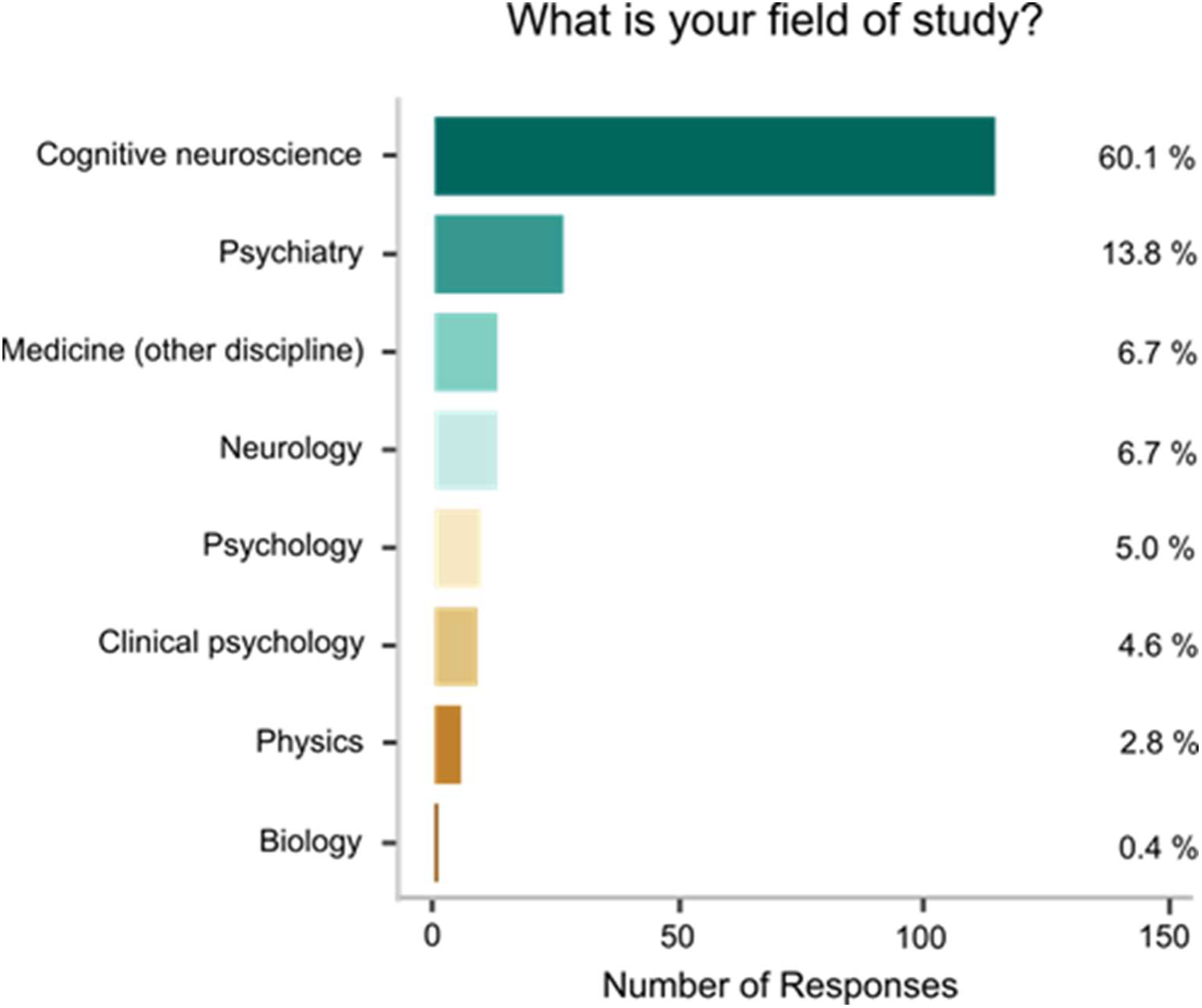
Field of study

**Figure 5.**
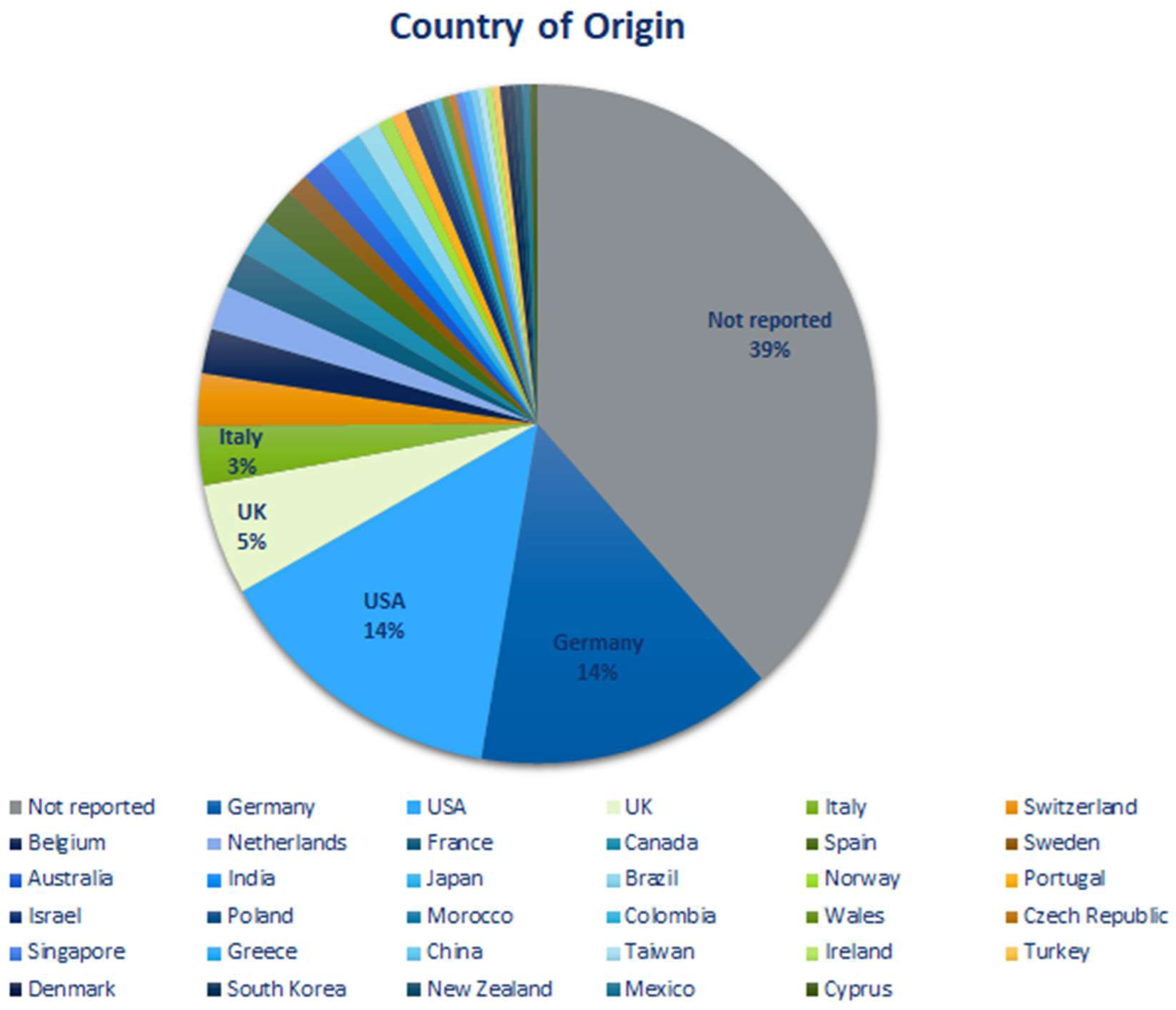
Country of residence

**Figure 6.**
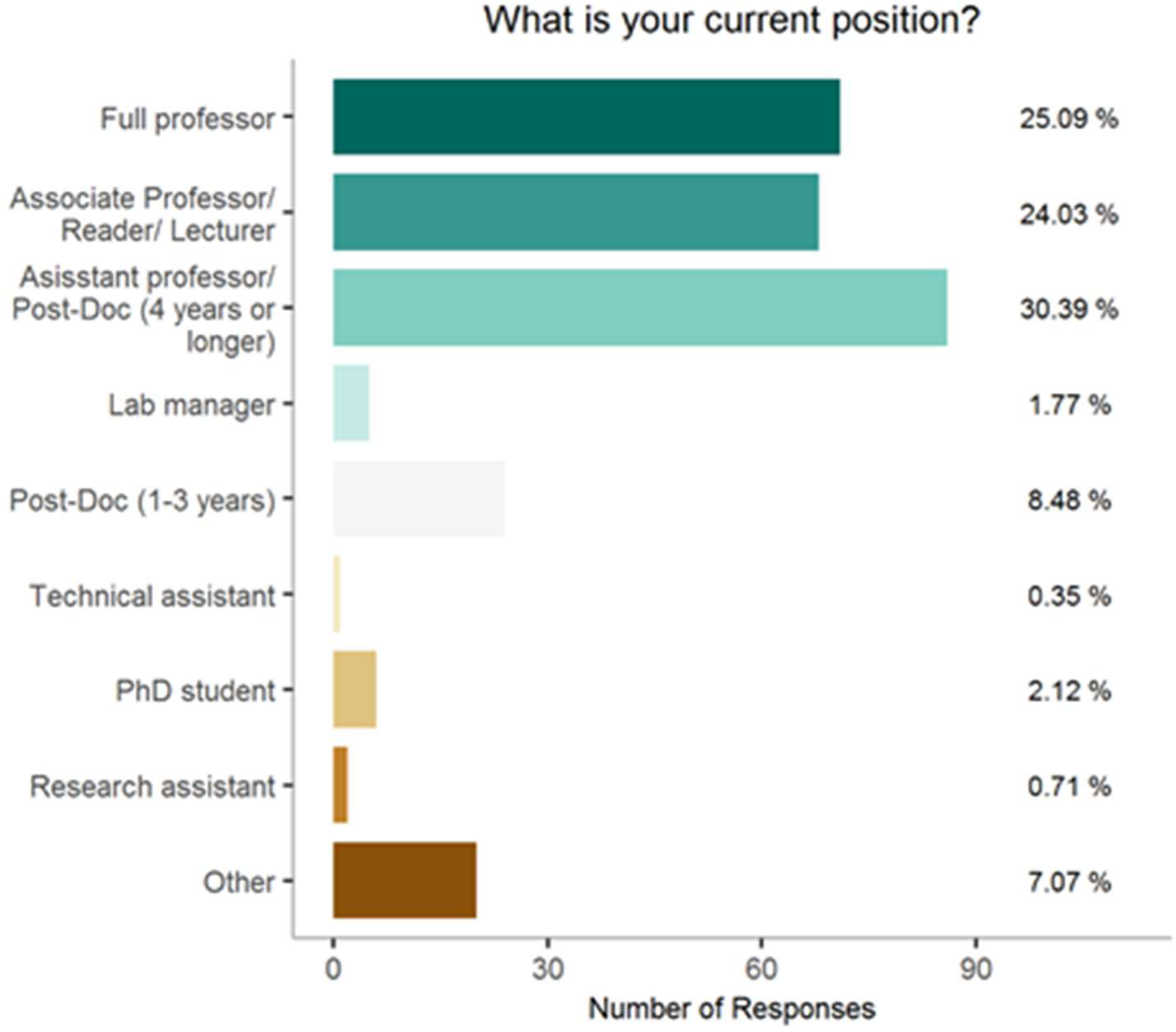
Job position and career level of the sample.

By clicking the personalized link, participants were navigated to an online form where they gave their informed consent before they could start with the questionnaire. This research was conducted in accordance with the declaration of Helsinki and was approved by the Ethics Committee of the Medical Faculty Mannheim of the University of Heidelberg.

### 2.2 Materials

The questionnaire was composed of five building blocks. Blocks 1-3 focused on three areas of open science practices: data structure, preregistration and data sharing. The fourth block asked about technical expertise with software (which was not analyzed for this publication) and the fifth part assessed sociodemographic data. In the beginning of each block a brief introduction to the topic area with definitions for key terms was provided. One or more questions on the subjective experience with the topic followed. Further, it included one or more questions to assess the likelihood to adopt practices of this topic area in the future on a 5-point Likert scale (“extremely unlikely” - 1, “somewhat unlikely” - 2, “neither likely nor unlikely” - 3, “somewhat likely” - 4, “extremely likely” - 5). The items for the data structure block were created by the author team with the major goal to assess knowledge and usage of BIDS in the fMRI community. Barriers and fears of adopting preregistration and data sharing practices were assessed by asking for agreement with statements on a 7-point Likert scale (“strongly disagree” - 1, “disagree” - 2, “somewhat disagree” - 3, “neither agree nor disagree” - 4, “somewhat agree” - 5, “agree” - 6, “strongly agree” - 7). For the data sharing block we used items from a previously published study on data sharing in psychology ^7^. Due to the broader scope of our survey and to reduce burden for participants, a selection of items and response options was drawn from Houtkoop et al. ^7^ and used in our questionnaire.

Furthermore, we restructured the item blocks from Houtkoop et al.’s survey. The items that Houtkoop et al. had grouped to a block on barriers related to data sharing were split up in one item block asking for preferred options to share data and in a second item block asking for a number of potential barriers. In the barriers-item block we merged these items with other items from Houtkoop et al.’s survey, which specifically assessed fear-related barriers of data sharing. The items on barriers for and fears of preregistration were inspired by the items on barriers for and fears of data sharing. For example, the preregistration item “preparing a preregistration is too time consuming for me” was based on the data sharing item “preparing data to make it suitable for online sharing is too time consuming for me”. Thus, several items from the preregistration block resembled items from the data sharing block which focused on comparable challenges such as lack of time, high complexity and lack of training in open science practices. Other items asked specific questions about each topic area (for example, “I am afraid that my preregistered hypotheses may turn out false” from the preregistration block or “I am afraid that other researchers will discover errors in my data’’ from the data sharing block). The online questionnaire was implemented using SoSci Survey ^30^.

### 2.3 Data analysis

Statistics software R version 4.0.5 was used to analyze the data. To analyze individual differences, we defined subgroups based on demographic variables of interest: 1) Career level (full/associate professors vs. assistant professors or lower stage), 2) years of research experience , 3) EU residency (EU resident vs. no EU resident) and 4) affiliation with medical faculty (university hospital/medical faculty vs. other faculty). T-tests were used to assess individual differences and Bayes Factor (BayesFactor Version 0.9.12-4.2 ^31^) was determined to assess the relative evidence for the alternative hypothesis versus the null hypothesis (BF10). We used the low information cauchy prior with a scale factor of 0.707, which is the default of the BayesFactor package that was used for this analysis and which has been suggested for psychological applications. Bayes factors take values between p(Data|H1) and p(Data|H0), with the common minimum cutoff of 3 (or below ⅓) indicating claims of evidence in favour of one hypothesis over the other. To explore latent variables that may drive responses to items on both data sharing and preregistration, an exploratory factor analysis was performed using R package lavaan_0.6-7 and psych_2.0.12 ^32^. An exploratory structural equation model was chosen to leverage the advantages of exploratory factor analysis and confirmatory factor analysis^33^, allowing the evaluation of exploratory models with goodness of fit measures. In total, the 28 statements that related to barriers and fears of data sharing and preregistration, as well as preference of how to share data, were used for the analyses. Each statement was rated on a Likert scale ranging from 1 (“strongly disagree”) to 7 (“strongly agree”). Factor analysis was performed using maximum likelihood estimation and oblique rotation (Oblimin), allowing factors to correlate with each other. The number of factors was determined using parallel analysis. Items with factor loadings >0.4 were retained.

To investigate whether groups with different response patterns exist, we performed a data-driven cluster analysis on the seven factors received from exploratory structural equation modeling. The euclidean distance was used to construct the dissimilarity matrix and clustering performed using Ward’s method. The optimal number of clusters was chosen based on the elbow and the silhouette method using the factoextra package version 1.0.7 ^34^. To explore whether any demographic variables could predict cluster belongingness, we performed a logistic regression with research experience, primary affiliation with medical faculty, EU residency, and career level as predictors. Model accuracy was calculated using the Caret package^35^.

## 3 Results

### 3.1 Preregistration is facing challenges

42.4 % participants indicated they have never preregistered a study. Among the rest of participants, the most frequently used preregistration platform was the Open Science Framework (OSF, 32.5%), followed by ClinicalTrials.gov (25.1%), and AsPredicted (9.5%). 14.1% indicated they had submitted a *registered report* article type ^36^ to a scientific journal (Figure 7). About the same number of participants who said they had preregistered a study before indicated they were likely or extremely likely to preregister their next study online (55%), while 26% disagreed (Figure 8). Asked about potential barriers for preregistration, 64% agreed at least to some extent with the statement that their analyses were too complex to preregister. The statement “There is no sufficient reward for preregistration” reached the second rank (53%). 46% agreed that preparing a preregistration is too time-consuming for them and 41% agreed that they know too little about preregistration platforms (41%) or that they have never learned to preregister a project (41%). 74% disagreed with the statement that they had never thought about preregistering a project (14% agreed). 10% indicated that their supervisor does not support preregistration. Asked about potential fears of preregistration, 49% agreed that they were afraid that their preregistered methods may turn out as suboptimal or inadequate. 23% agreed they were afraid that their preregistered hypotheses may turn out false. We also asked whether participants think that it is necessary to register studies with an explorative research question and 48% agreed (Figure 9).

**Figure 7.**
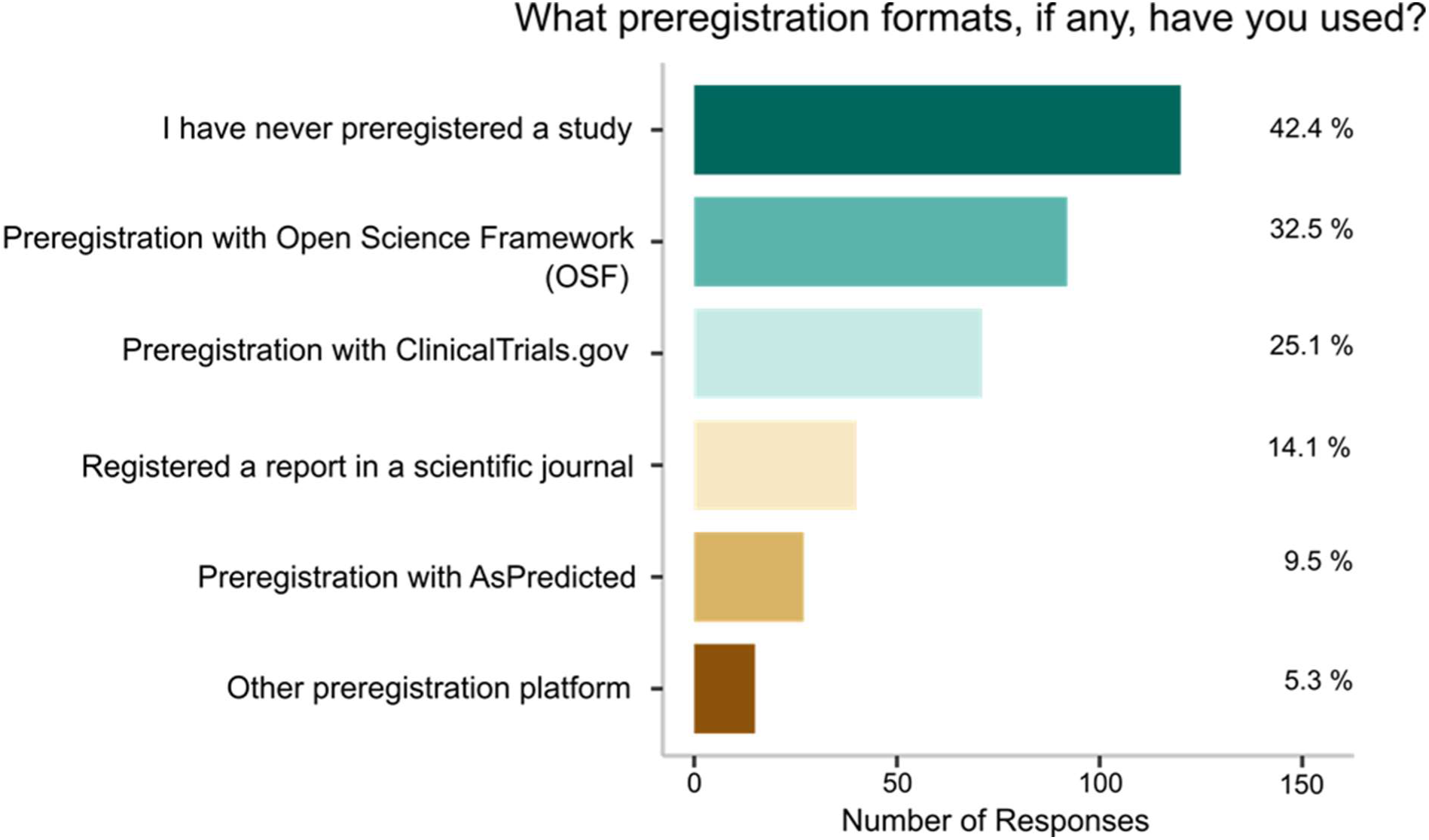
Preregistration formats used in the past.

**Figure 8.**
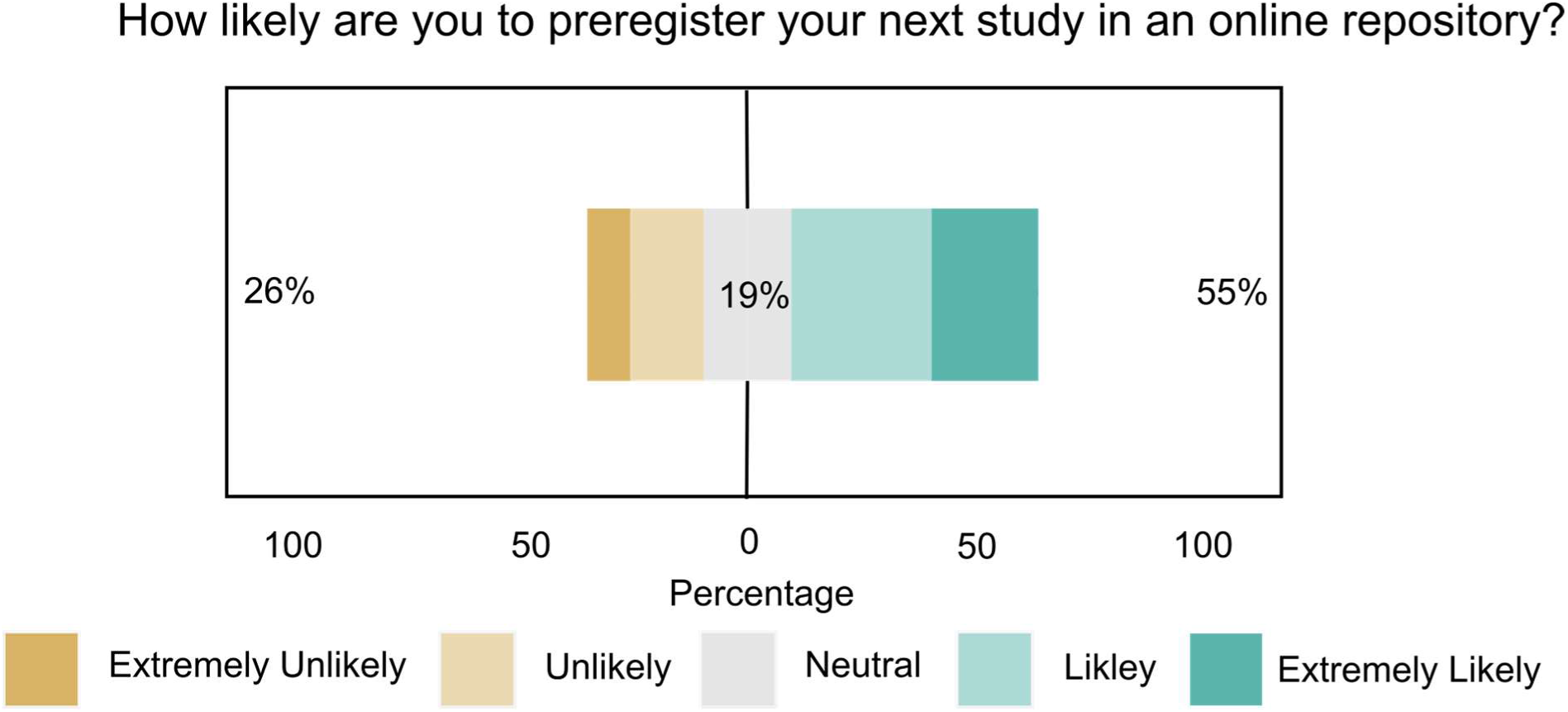
Intention to preregister in the future.

**Figure 9.**
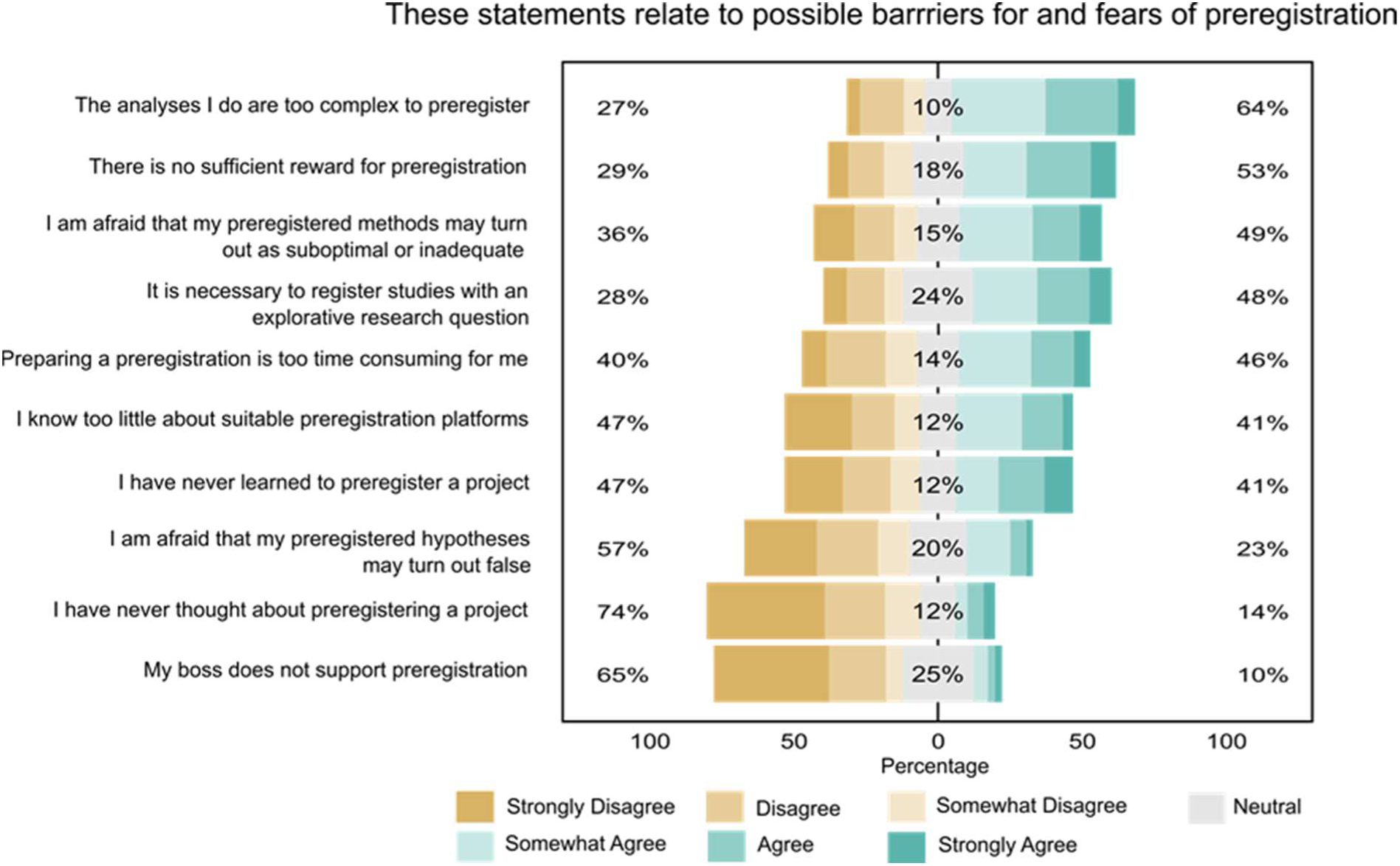
Potential barriers for and fears of preregistration.

### 3.2 Sharing raw data is common practice for many

66% of all participants said they have shared neuroimaging raw data with other researchers outside their department before. Asked about the intention to share primary research data of their next neuroimaging paper in an online repository, 54% indicated they were likely or extremely likely to do this, while 25% were unlikely or extremely unlikely (Figure 10). Asked whether they were not allowed to share primary neuroimaging data due to legal constraints, 64% disagreed at least to some extent, while 9% agreed (27% did neither agree nor disagree, Figure 11). If a participant did not disagree strongly with the above statement, a follow up question was asked to investigate the reasons why the participant thought s/he was not allowed to share primary neuroimaging research data. Most participants endorsed the statement that anonymity cannot be guaranteed if the data is shared (45.2% agreed at least somewhat). 41% indicated their consent forms state that data will not be shared. 29.5% responded that their institutional review board does not allow them to share data. 14.8% reported stakeholder interests prohibiting data from being shared and 6.7% said that a funder, advisor or supervisor does not allow them to share data (Figure 12).

**Figure 10.**
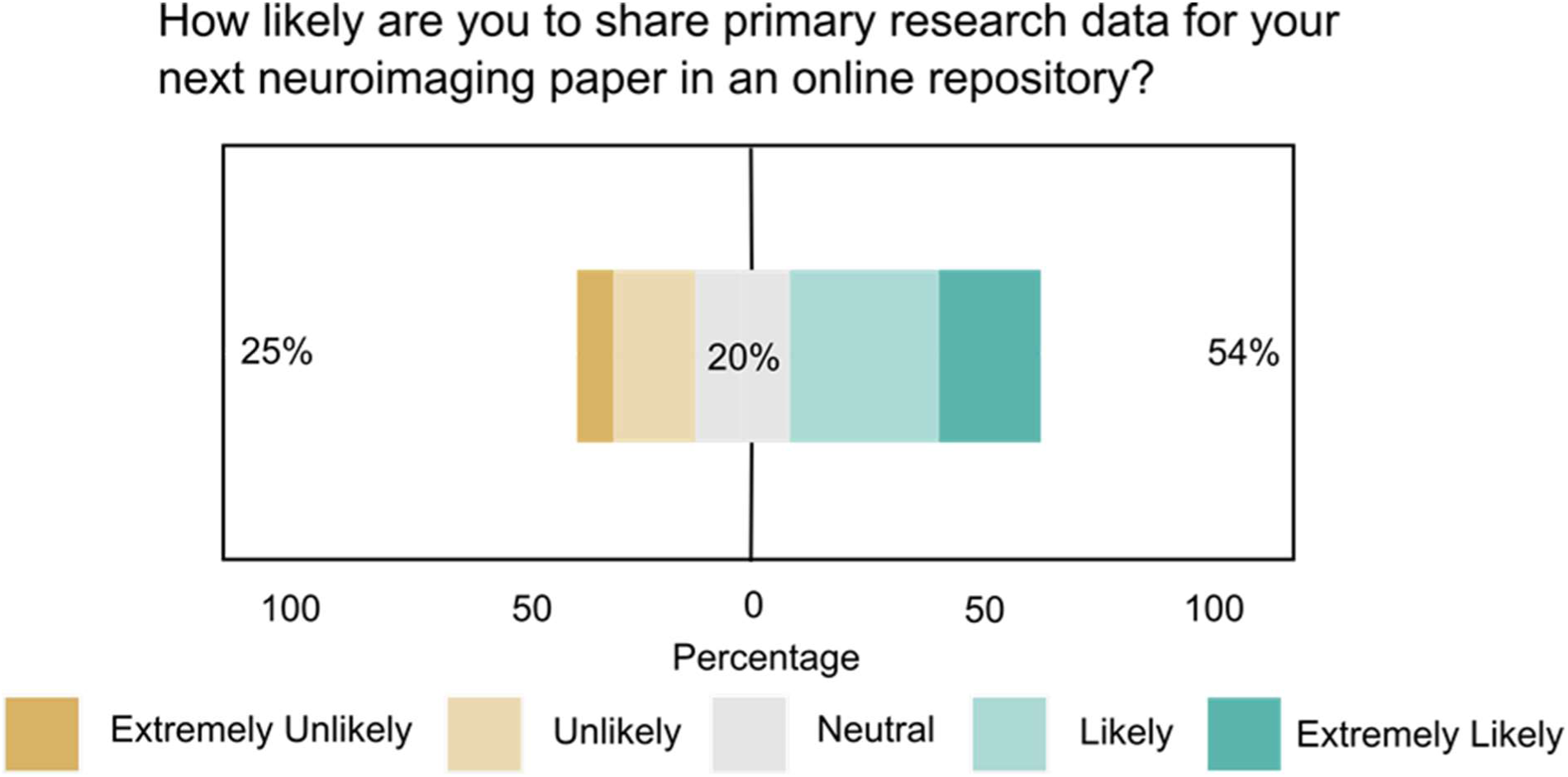
Intention to share data for the next neuroimaging paper.

**Figure 11.**
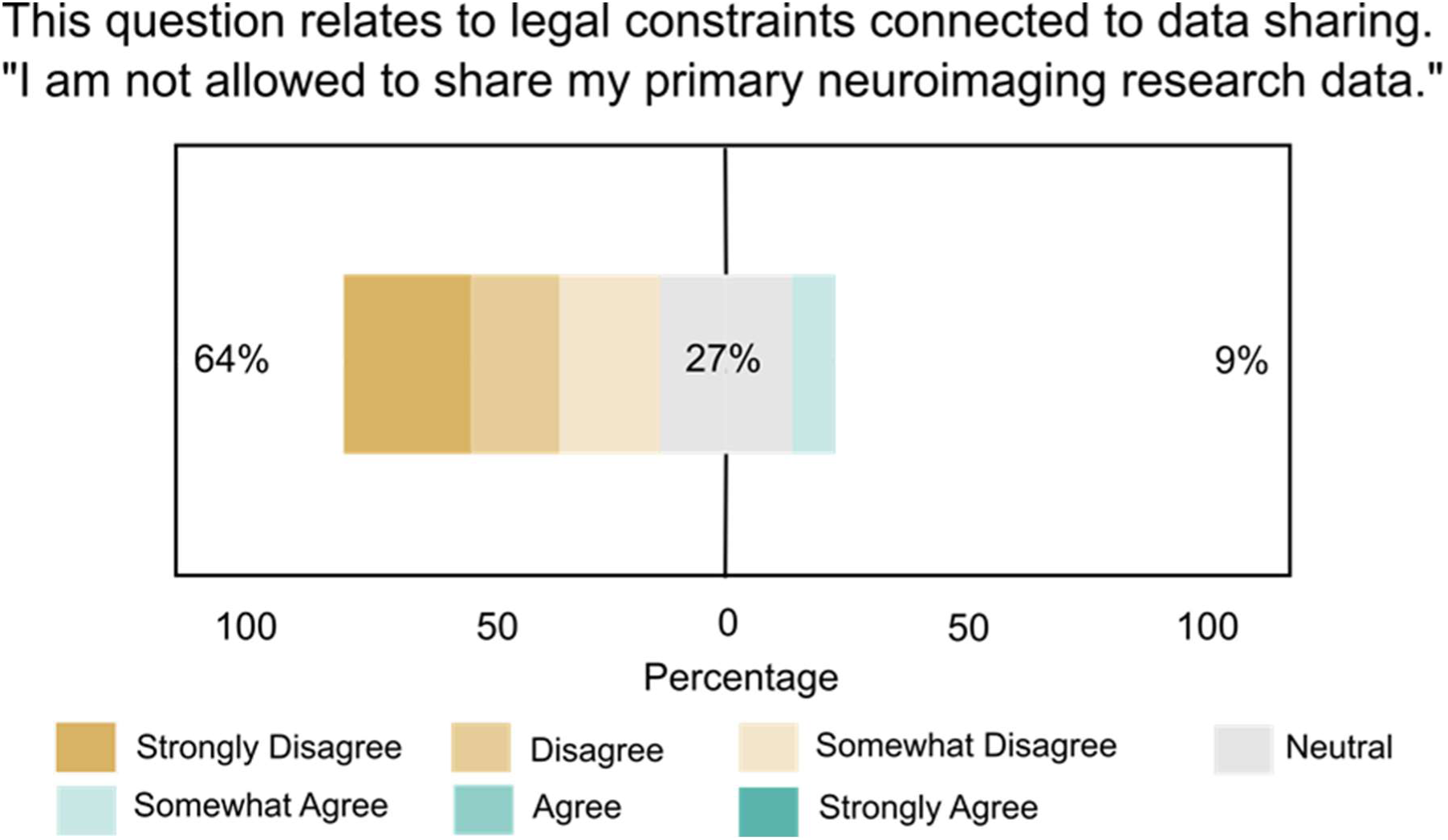
Not allowed to share primary data due to legal constraints.

**Figure 12.**
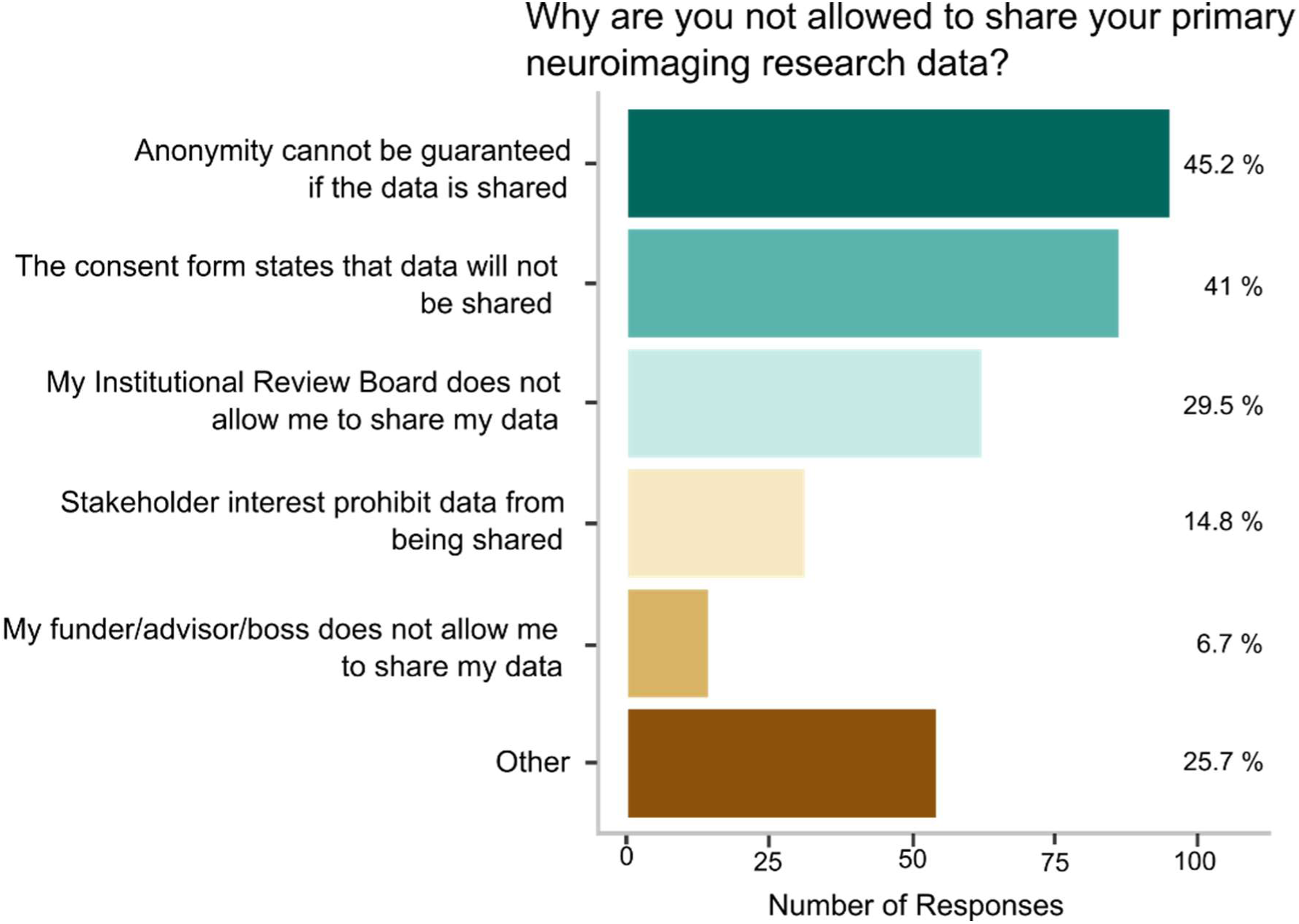
Why not allowed to share primary neuroimaging data.

### 3.3 Europeans more hesitant to share raw data online in the future

To explore interindividual differences that may result from national data protection legislation, we compared participants who indicated their country of residence within the European Union (EU) vs. outside the EU. The number of participants who indicated they had shared data in the past outside their department did not significantly differ between EU and non EU researchers (*Χ*^2^(1)= 0.287, p = 0.591). More participants from the EU agreed with the statement they are not allowed to share primary neuroimaging data for legal reasons, t(251.94) = 2.84, p<0.005, BF10=6.26, and less participants from the EU agreed they will likely share primary research data from their next neuroimaging paper online, t(269.59) = 3.09, p<0.002, BF10=10.75.

### 3.4 Researchers appreciate data sharing agreements

To learn more about the preferred mode of data sharing, we let participants evaluate several options on how data can be shared with other researchers. Highest agreement was found for the option to share data under a data sharing agreement to be signed by the recipient (65%), directly followed by the option to share upon personal request and therewith bypassing a data repository (64%). With 58% agreement, sharing via a managed online repository with restricted access found high approval, too. The option to share via an online repository with unrestricted access was prefered by 35% of participants, while 45% expressed disagreement with this item. 17% prefered that researchers with reasonable interest can work with their data, but that this work needs to be done on the server of their home institution (63% disagreed). Finally, 6% agreed they preferred not to give away raw data to other researchers, whereas 81% disagreed (Figure 13).

**Figure 13.**
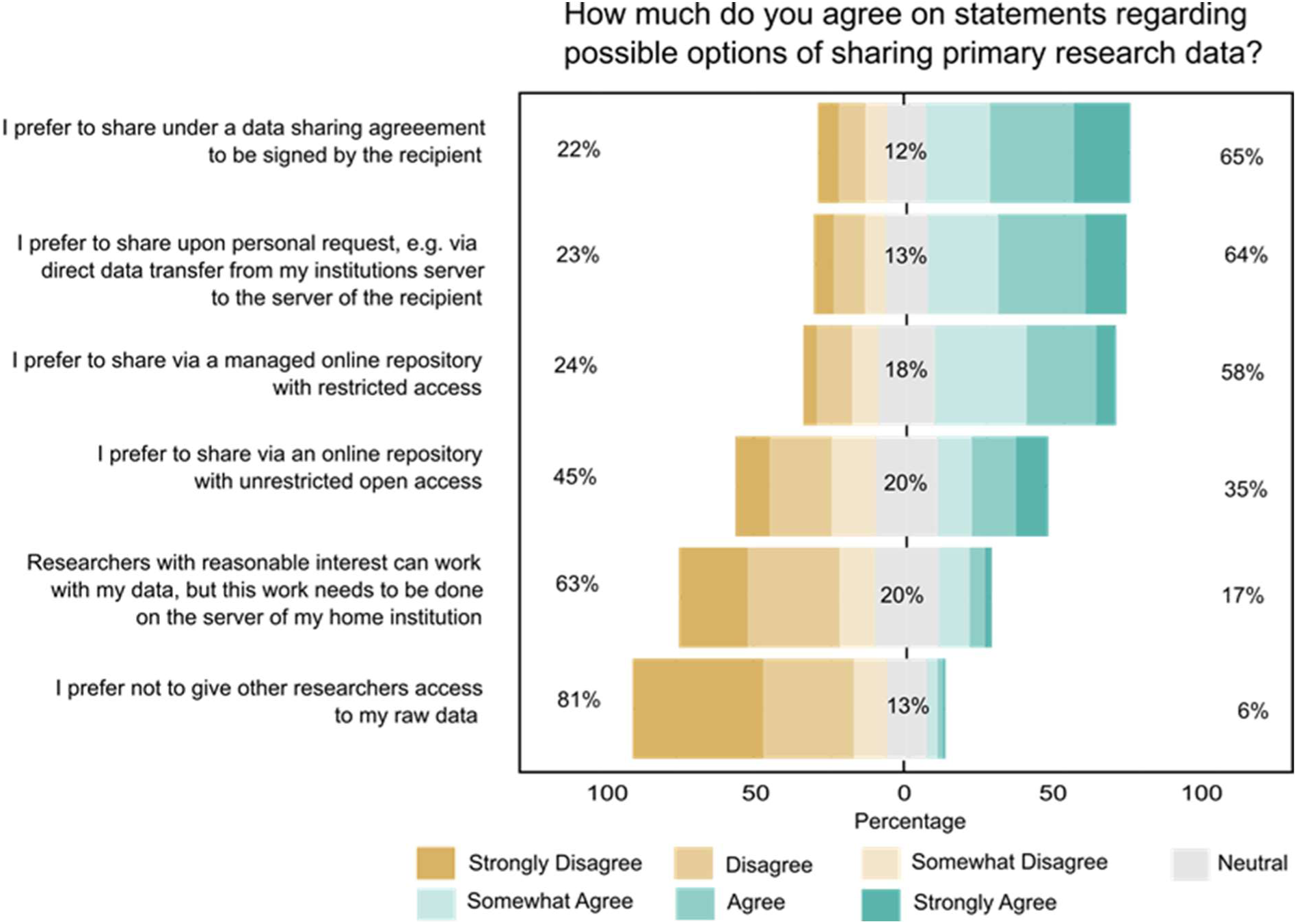
Possible options of data sharing.

### 3.5 Lack of resources poses a high hurdle to data sharing

Asked about barriers for and fears of data sharing, 67% agreed at least somewhat that preparing data to make it suitable for online sharing is too time-consuming. The second leading statement “I lack funding to make data suitable for online sharing” received 61% agreement. 47% of participants agreed they are afraid of being scooped, i.e., that other researchers may publish results received with their data set before they can. 41% agreed they knew too little about suitable data repositories and 40% agreed they never learned to share their research data online. 38% endorsed the statement they are afraid not to get proper recognition for sharing data. The concern that data sets were too big (33%) or too complex (30%) to share were found on the following ranks. 25% expressed fears that other researchers could run alternative analyses on their data to rebut their own conclusions and 24% agreed they are afraid that other researchers will discover errors in their data. 11% agreed their supervisor does not support online data sharing. 11% agreed they have never thought about data sharing, whereas 81% disagreed (Figure 14).

**Figure 14.**
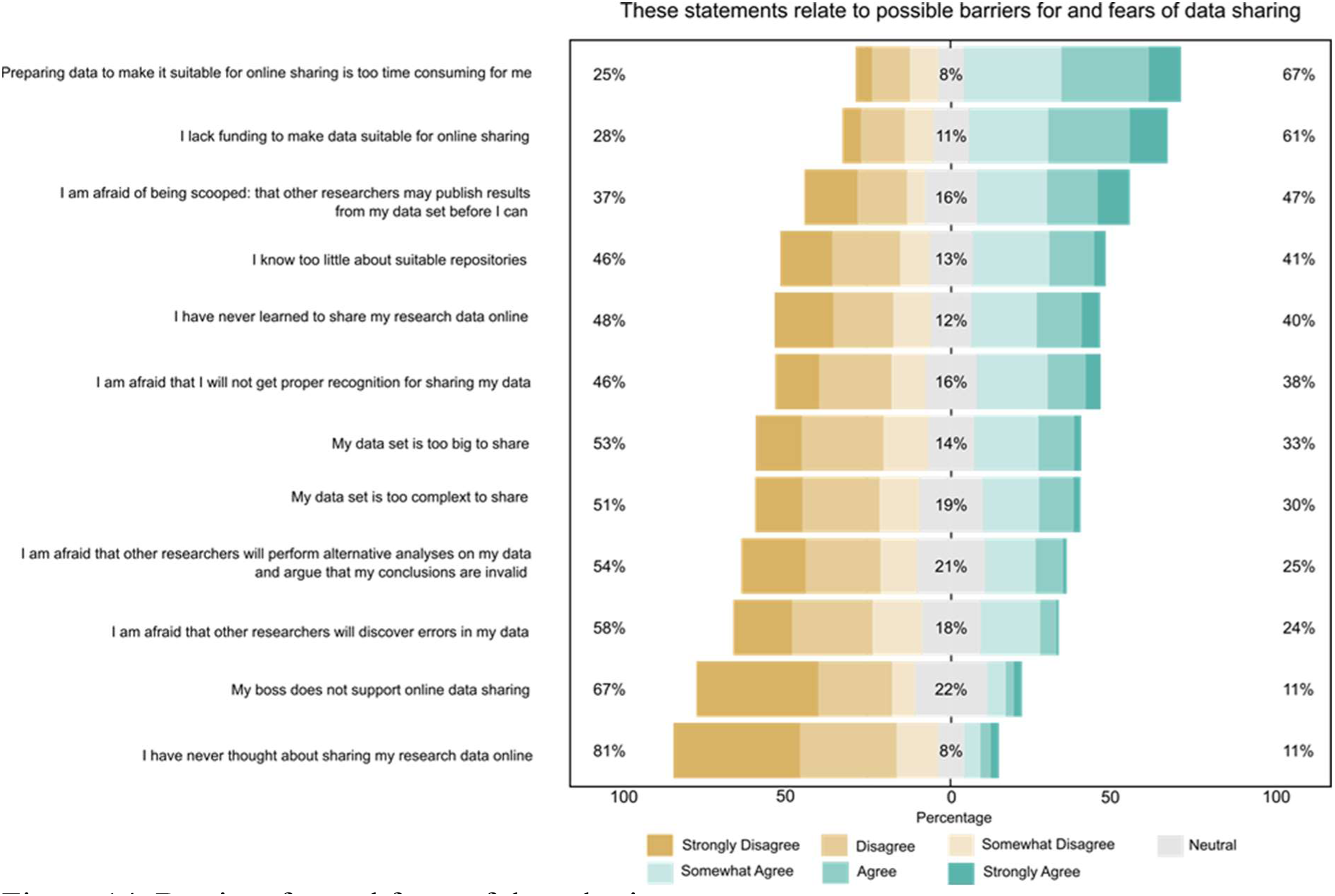
Barriers for and fears of data sharing.

### 3.6 High interest in using BIDS

72% of respondents indicated that they had heard about BIDS before. 35% said that they had used BIDS in the past and have been working with it for 2.27 (1.78) years on average. The vast majority, 91%, find it likely or extremely likely that they are going to use BIDS in the future (Figure 15). Participants who said that they have not used BIDS before were asked to report the reason. Most indicated they had not heard about BIDS before (41.5%), they had no time to implement it in the lab (36.1%), or to learn more about it (28.4%). 12.6% agreed they were lacking technical expertise to get BIDS conversion running, 10.9% said they were currently implementing it, and 6% said they were using a different data structure format than BIDS. 5.5% deemed BIDS not relevant for their lab (Figure 16). Those preferring to operate software via graphical user interface (GUI) used BIDS significantly less often as compared participants who prefer to interact via command interface, *Χ*^2^ (1)= 18.72 , p < 0.001. Those who indicated that they had used BIDS before were then asked about experience with BIDS-compatible software: 32% participants experienced with BIDS used custom code to convert raw neuroimaging data into the BIDS format, while 16% indicated that they have not used any conversion software (Figure 17). Several participants confirmed they have been using software that can operate on BIDS formatted data sets such as fMRIPrep ^20^ (44%), MRIQC ^37^ (23%), OpenNeuro ^38^ (18%) and other tools (<10%) (Figure 18).

**Figure 15.**
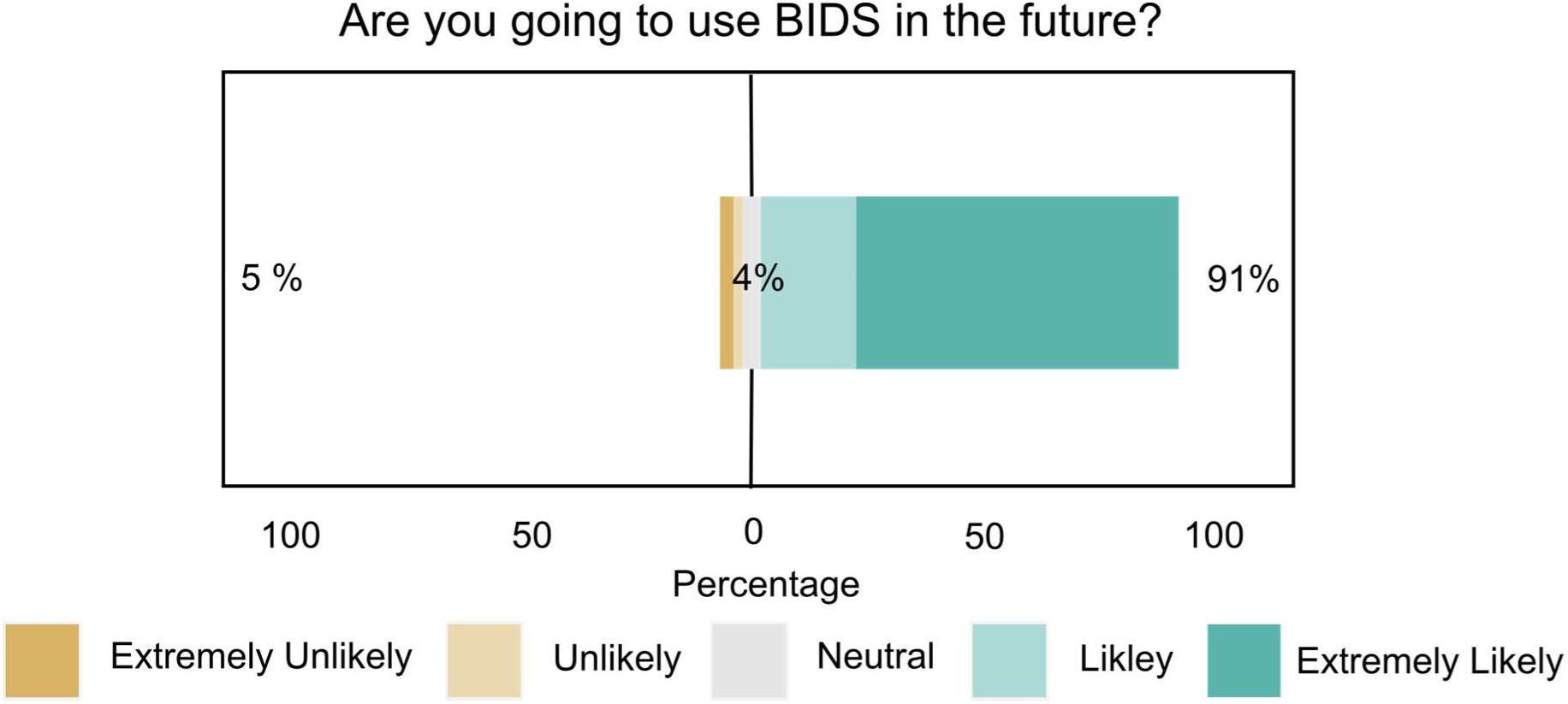
Intention of using BIDS.

**Figure 16.**
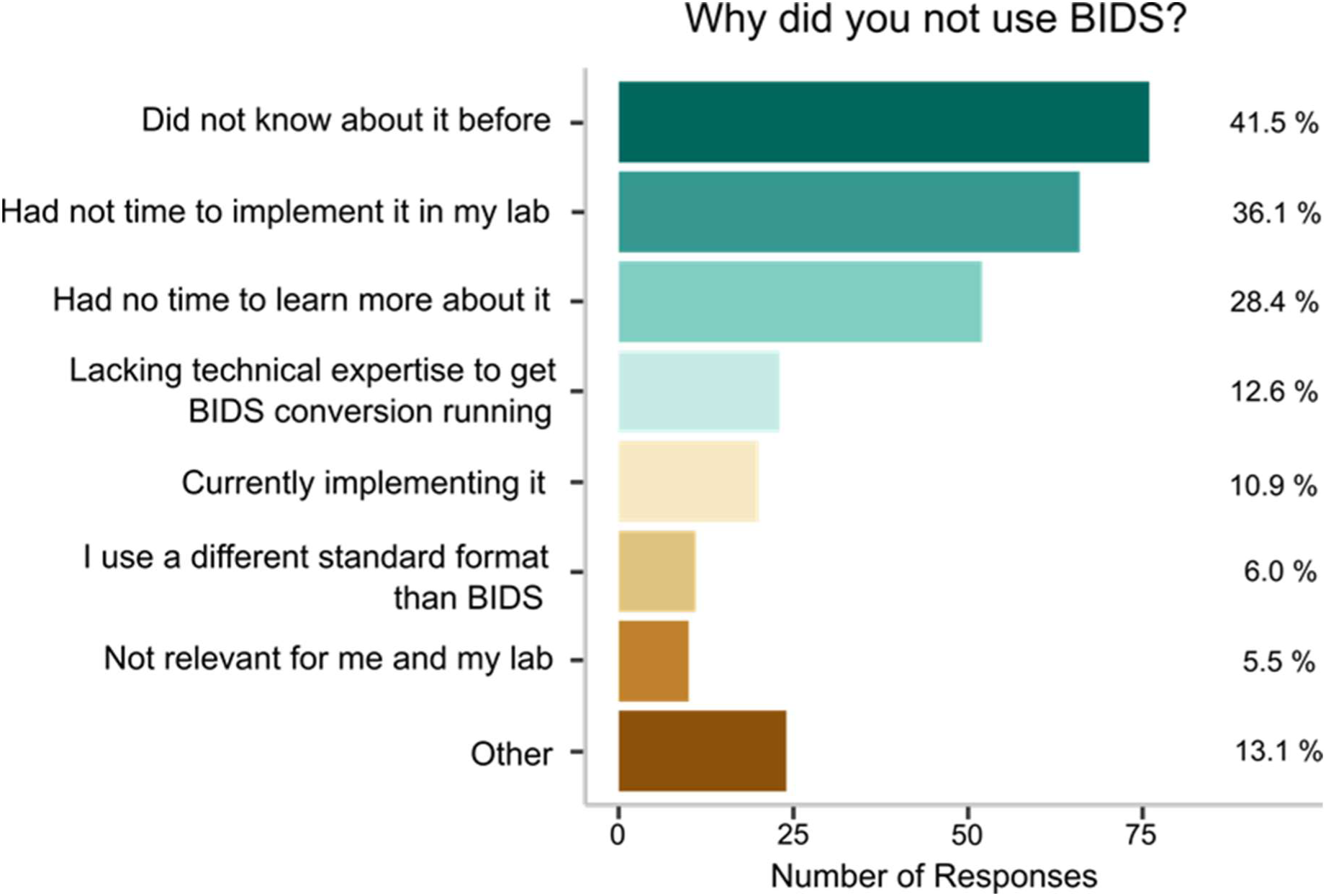
N=183 participants have not used BIDS before and were asked why. Participants could check one or more response options. Bars show the percentage of responses per option.

**Figure 17.**
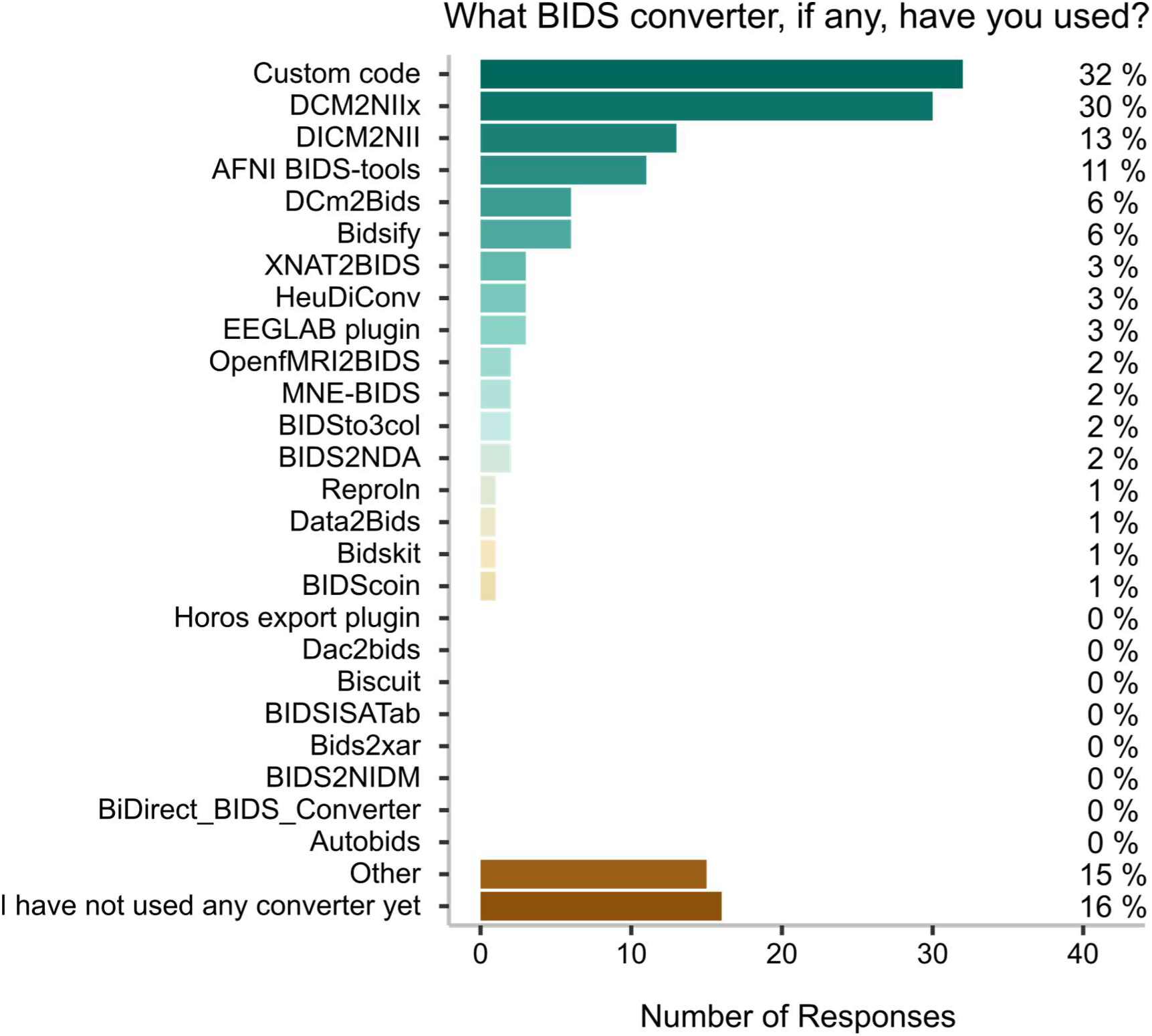
N=101 participants have used BIDS before and were asked what conversion software they had used. Participants could check one or more response options. Bars show the percentage of responses per option.

**Figure 18.**
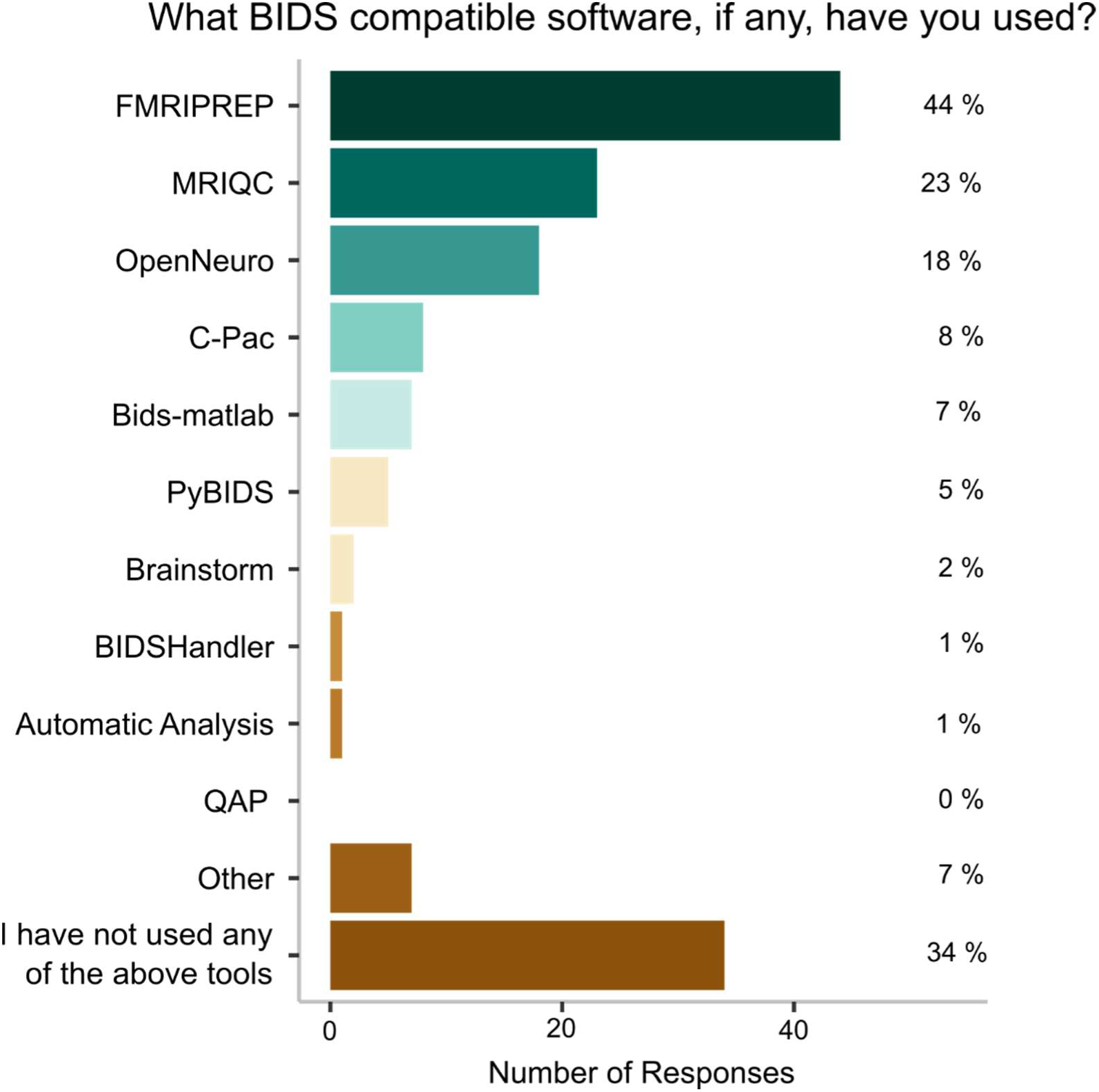
N=101 participants have used BIDS before and were asked what BIDS compatible software they had used. Participants could check one or more response options. ars show the percentage of responses per option.

### 3.7 Factors underlying barriers, fears, and preferences of preregistration and data sharing

We explored whether the answers of our participants could be reduced to a smaller set of interpretable latent variables. Bartlett’s test confirmed that the items correlated sufficiently, *Χ*² (378) = 3135.5, *p*<0.001, to explore the structure with factor analytic methods. The KMO test indicated overall acceptable Measure of Sample Adequacy (MSA=0.81). On item level, the MSA suggested the inadequacy of the item “It is necessary to register studies with an explorative research question” (MSA=0.46). We excluded the item, due to the low MSA and as it does not name a barrier or fear as the rest of the items. While parallel analysis recommended the eight-factor solution, we decided to choose a seven-factor model, due to parsimony of this solution, as it already provided good model fit (Table 2): The Comparative Fit Index (CFI) reached 0.937 (cut off >0.9) while the Root Mean Square Error of Approximation (RMSEA) was below the cut-off of 0.05 (RMSEA=0.042). The seven factors resulting from this analysis included: fear of being transparent, lack of experience regarding preregistration, lack of experience regarding data sharing, complexity of own research, need for data governance, unsupportive environment, and lack of resources for data sharing.

**Table 1.**
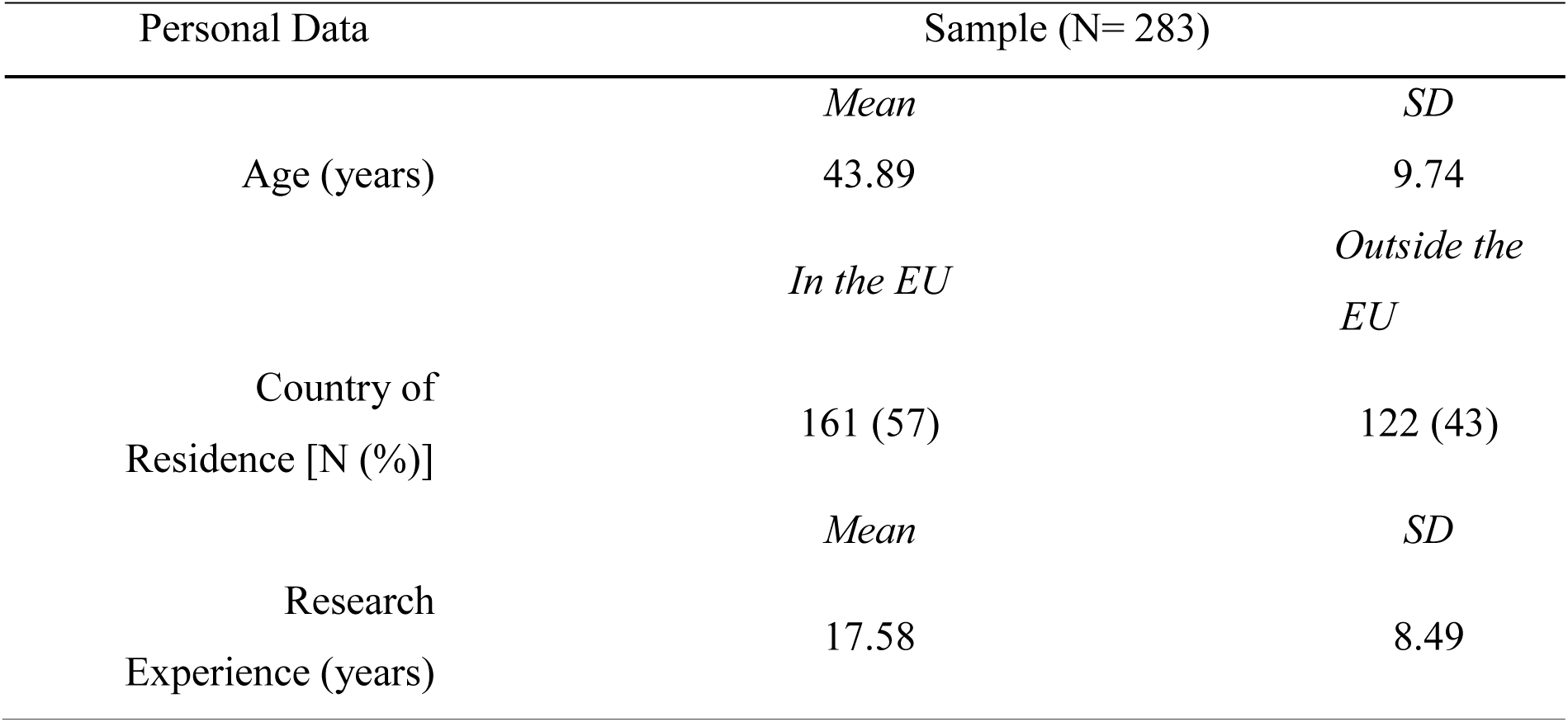
Sample characteristics

**Table 2.**
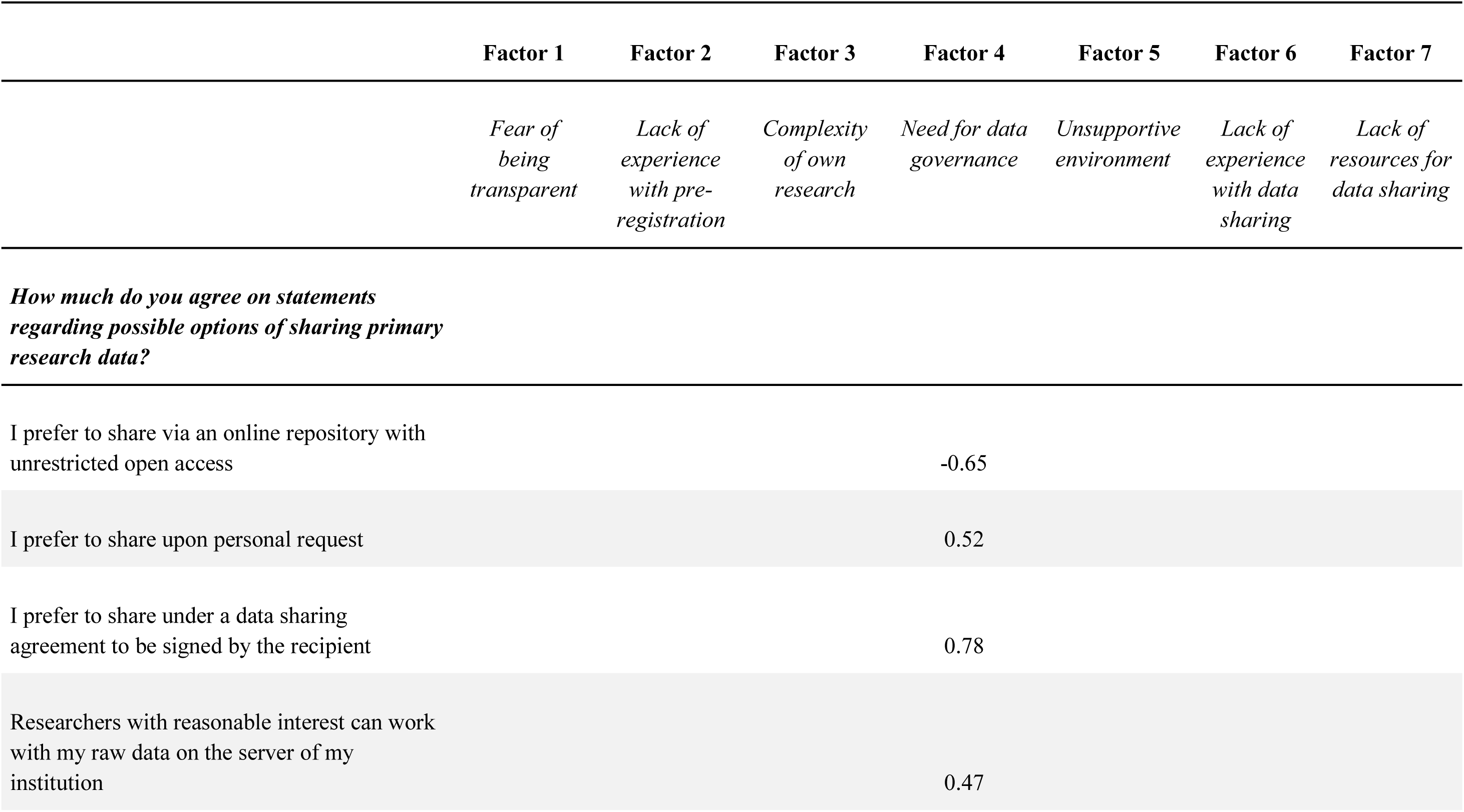
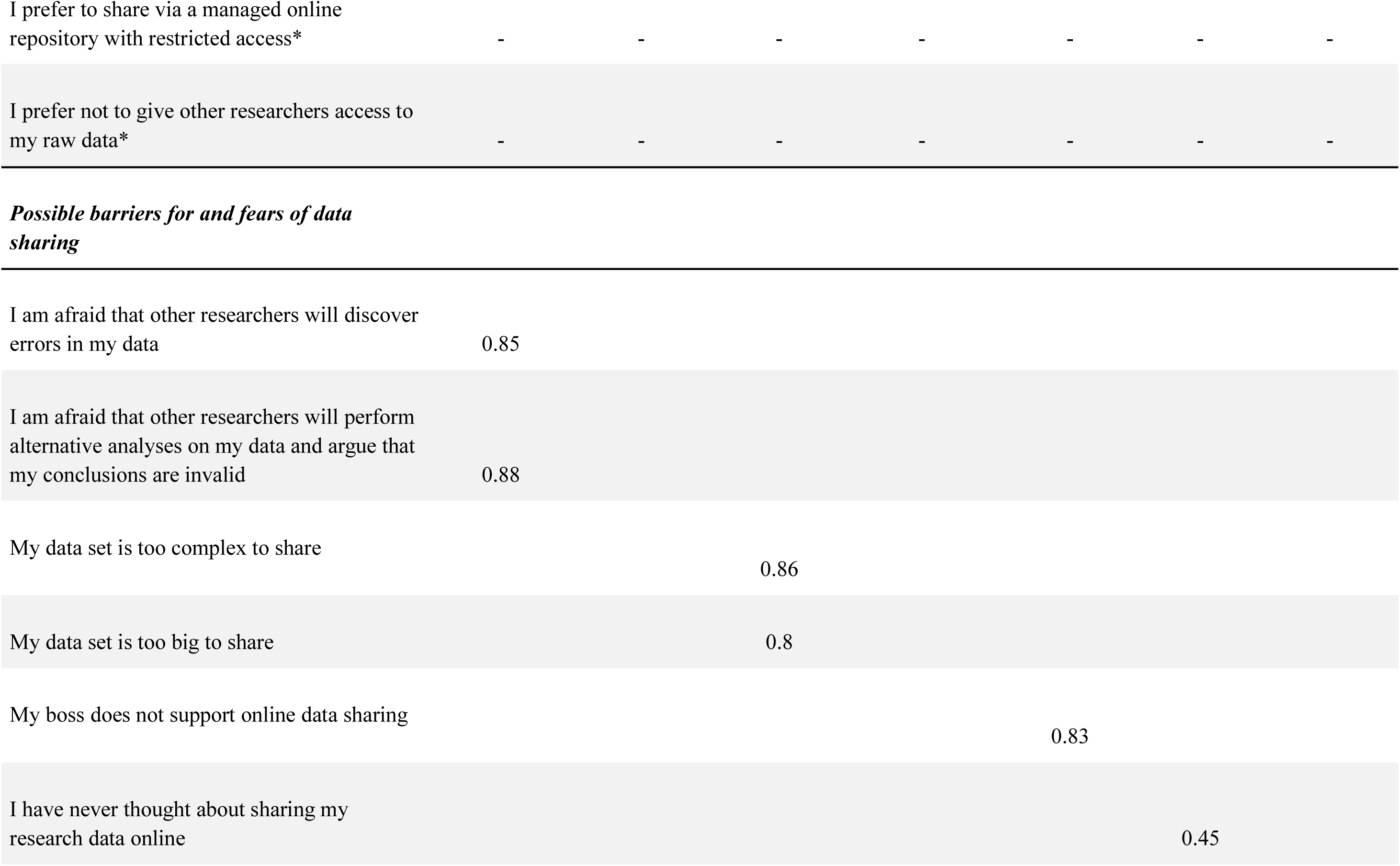
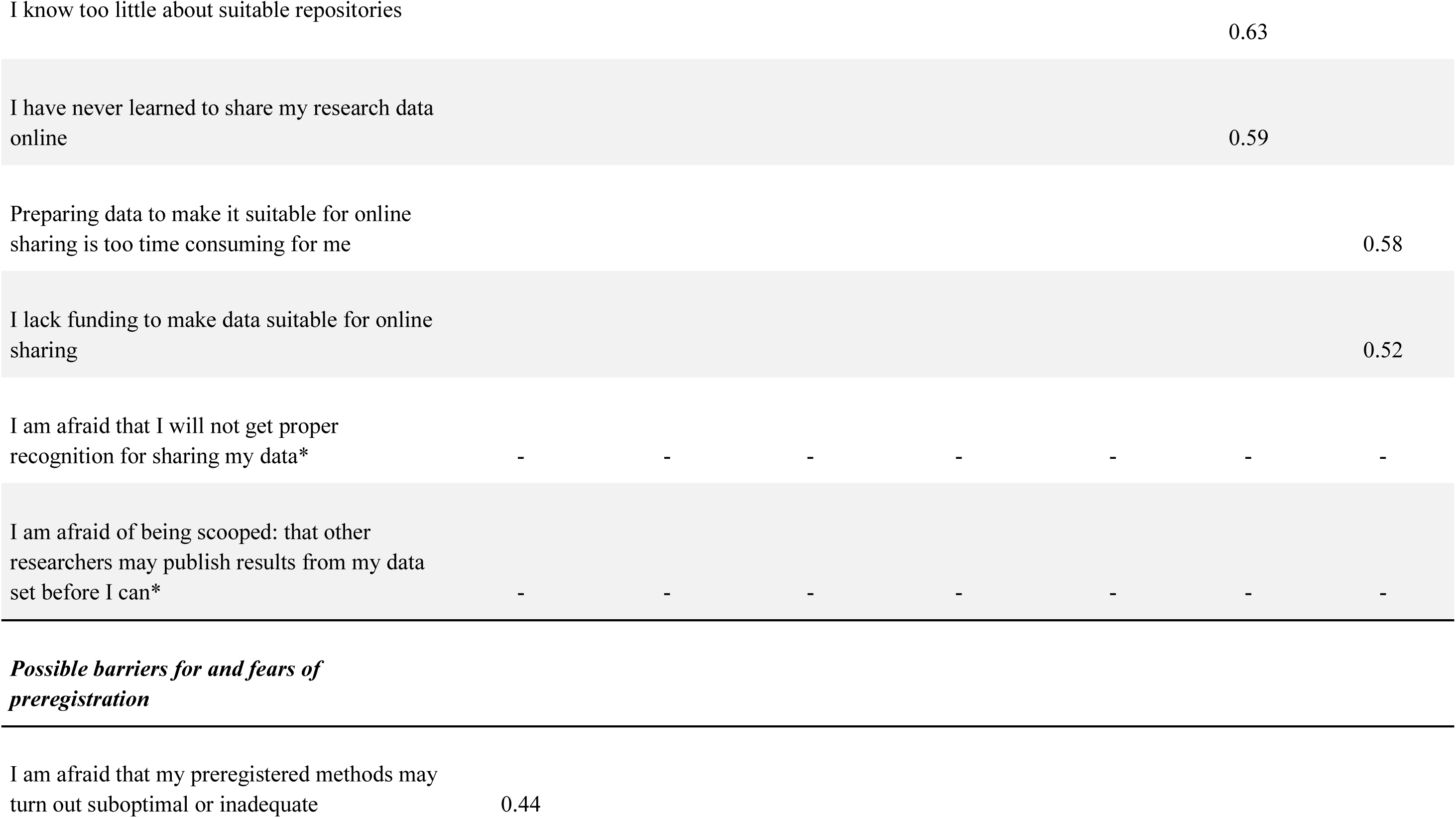
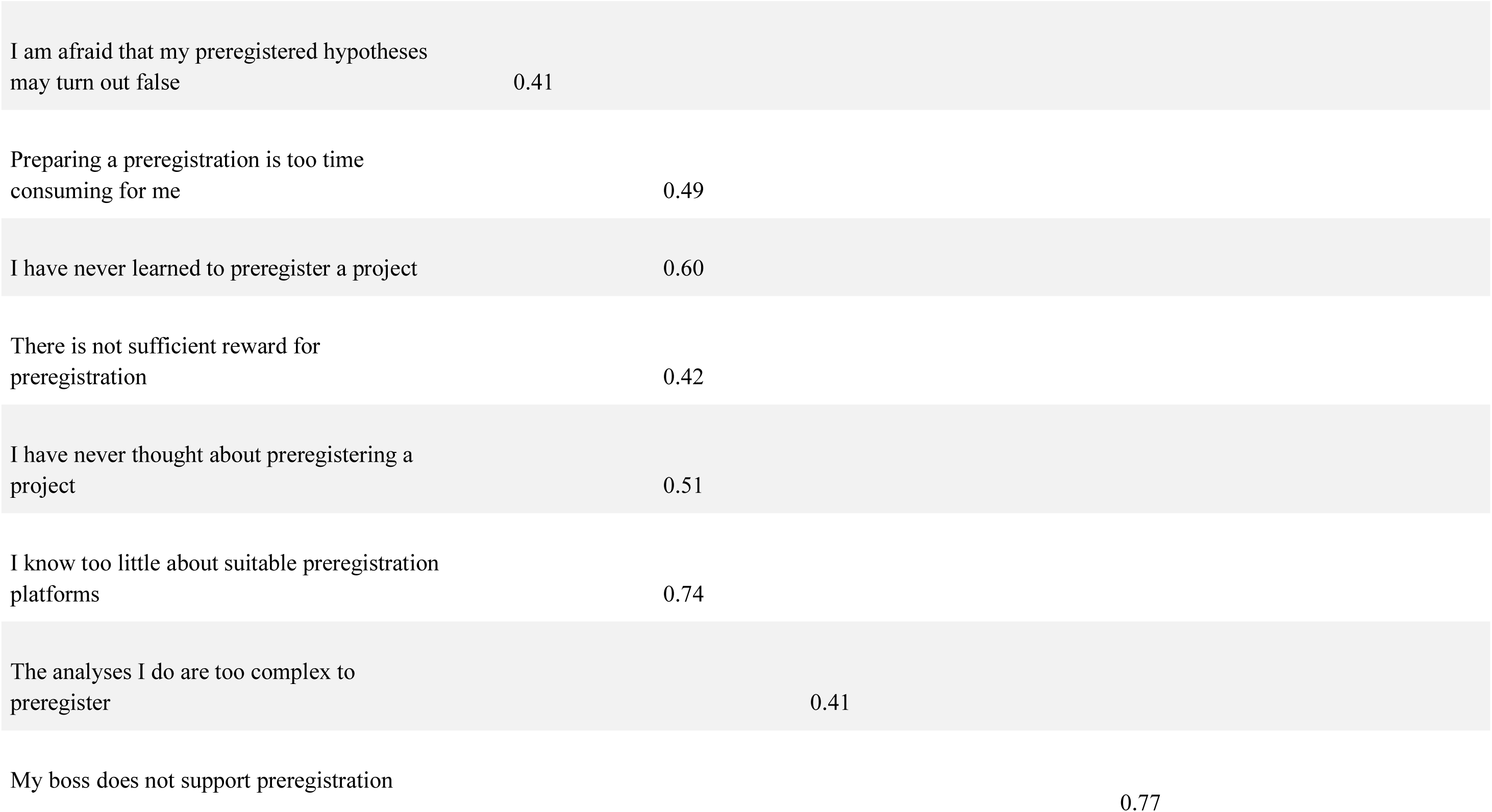
. Results from Exploratory Structural Equation

We used the results from factor analysis to build seven subscales from our questionnaire. For each participant we calculated subscale scores by averaging the item scores assigned to each factor. The subscale scores were further used to explore individual differences, comparing participants based on demographic variables. The Bonferroni corrected results of all performed comparisons can be found in Table 3. For the factor “fear of being transparent” we found that people with a lower career level were significantly more fearful than people with a higher career level. For “need for data governance”, people having their primary affiliation with a medical faculty showed significantly higher scores than people having their primary affiliation with a psychological or other faculty. Respondents residing in the EU had a higher need for data governance than non-EU residents. Lastly, people affiliated with a medical faculty scored higher on “unsupportive environment”, as did respondents with a lower career level compared to respondents with a higher career level.

**Table 3.** Results of significance testing for the demographic variables “Research experience”, “EU residency” , “Primary affiliation” and “Career level”. Results are significant at a corrected p<0.0017 using Bonferroni correction.

|  | Research experience |  |  |  |  |  |
| --- | --- | --- | --- | --- | --- | --- |
|  | <16 years<br>experience | >16 years<br>experience | *t*-test results |  |  |  |
|  | (*n*=145) | (*n*=138) |  |  |  |  |
|  | *mean* | *mean* | *t* | *df* | *p* | *BF*<sub>10</sub> |
| Fear of being transparent | 3.49 | 3.19 | 1.89 | 278.83 | 0.06 | 0.72 |
| Lack of experience |  |  |  |  |  |  |
| preregistration | 3.65 | 3.81 | -1.04 | 277.28 | 0.30 | 0.22 |
| Complexity of |  |  |  |  |  |  |
| own research | 3.43 | 3.76 | -2.02 | 280.29 | 0.04 | 0.90 |
| Need for data governance | 4.06 | 4.26 | -1.30 | 281.00 | 0.19 | 0.29 |
| Unsupportive environment | 2.74 | 2.30 | 2.53 | 280.90 | 0,012 | 2.67 |
| Lack of experience |  |  |  |  |  |  |
| data sharing | 3.29 | 3.08 | 1.17 | 280.02 | 0.24 | 0.25 |
| Lack of resources |  |  |  |  |  |  |
| for data sharing | 4.41 | 4.84 | -2.40 | 280.85 | 0,016 | 1.98 |

|  | Primary affiliation |  |  |  |  |  |
| --- | --- | --- | --- | --- | --- | --- |
|  | ...with<br>university<br>hospital/medical<br>faculty | ...with<br>psychological<br>or other<br>faculty | *t*-test results |  |  |  |
|  | (*n*=139) | (*n*=144) |  |  |  |  |
|  | *mean* | *mean* | *t* | *df* | *p* | *BF*<sub>10</sub> |
| Fear of being transparent | 3.46 | 3.23 | 1.29 | 244.90 | 0.20 | 0.31 |
| Lack of experience |  |  |  |  |  |  |
| preregistration | 3.85 | 3.60 | 1.56 | 244.80 | 0.12 | 0.44 |
| Complexity of |  |  |  |  |  |  |
| own research | 3.65 | 3.58 | 0.40 | 242.92 | 0.69 | 0.15 |
| Need for data governance | 4.78 | 3.81 | 4.21 | 234.14 | *<0.001 | 496.00 |
| Unsupportive environment | 2.87 | 2.15 | 4.04 | 230.99 | *<0.001 | 264.04 |
| Lack of experience |  |  |  |  |  |  |
| data sharing | 3.37 | 2.99 | 2.05 | 242.60 | 0.0415 | 1.01 |
| Lack of resources |  |  |  |  |  |  |
| for data sharing | 4.81 | 4.46 | 1.80 | 243.35 | 0.07 | 0.64 |

|  | EU residency |  |  |  |  |  |
| --- | --- | --- | --- | --- | --- | --- |
|  | Yes | No | *t*-test results |  |  |  |
|  | (*n*=161) | (*n*=122) | for EU residency |  |  |  |
|  | *mean* | *mean* | *t* | *df* | *p* | *BF*<sub>10</sub> |
| Fear of being transparent | 3.38 | 3.31 | 0.43 | 267.13 | 0.67 | 0.14 |
| Lack of experience |  |  |  |  |  |  |
| preregistration | 3.66 | 3.81 | -0.89 | 243.20 | 0.37 | 0.20 |
| Complexity of |  |  |  |  |  |  |
| own research | 3.57 | 3.63 | -0.38 | 267.10 | 0.70 | 0.14 |
| Need for data governance | 4.36 | 3.89 | 3.09 | 250.66 | *<0.001 | 12.65 |
| Unsupportive environment | 2.70 | 2.42 | 1.07 | 272.81 | 0.28 | 0.22 |
| Lack of experience |  |  |  |  |  |  |
| data sharing | 3.25 | 3.09 | 0.91 | 260.46 | 0.36 | 0.20 |
| Lack of resources |  |  |  |  |  |  |
| for data sharing | 4.56 | 4.69 | -0.72 | 245.63 | 0.47 | 0.17 |

|  | Career level |  |  |  |  |  |
| --- | --- | --- | --- | --- | --- | --- |
|  | Full/Associate<br>Professor | Other | *t*-test results |  |  |  |
|  | (*n*=123) | (*n*=124) | for career level |  |  |  |
|  | *mean* | *mean* | *t* | *df* | *p* | *BF*<sub>10</sub> |
| Fear of being transparent | 3.02 | 3.66 | -4.23 | 280.16 | *<0.001 | 545.9 |
| Lack of experience |  |  |  |  |  |  |
| preregistration | 3.60 | 3.84 | -1.56 | 280.87 | 0.12 | 0.42 |
| Complexity of |  |  |  |  |  |  |
| own research | 3.59 | 3.59 | -0.03 | 272.55 | 0.98 | 0.13 |
| Need for data governance | 4.22 | 4.10 | 0.79 | 279.79 | 0.43 | 0.18 |
| Unsupportive environment | 2.22 | 2.83 | -3.56 | 276.69 | *<0.001 | 47.45 |
| Lack of experience |  |  |  |  |  |  |
| data sharing | 2.97 | 3.39 | 2.39 | 281.00 | 0,017 | 1.93 |
| Lack of resources |  |  |  |  |  |  |
| for data sharing | 4.73 | 4.51 | 1.24 | 279.37 | 0.22 | 0.27 |

### 3.8 Distinct subgroups of open science profiles

We explored whether there are groups of participants with distinct profiles, according to scores achieved on the subscales, which might serve as potential target groups for future actions on open science practices. The suggested optimal number of clusters was two, which was supported by the highest Dunn Index for the two-cluster solution (0.155), compared to the three- and four-cluster solutions (Figure 19). As visible in the profile plot (Figure 20), cluster 1 consists of researchers with less experience, more complex datasets, and more concerns regarding data sharing and preregistration, as well as a less supportive environment and fewer resources for data sharing. Cluster 2 was composed of researchers who were more experienced with open science practices and who saw overall less barriers and had lower fears.

**Figure 19.**
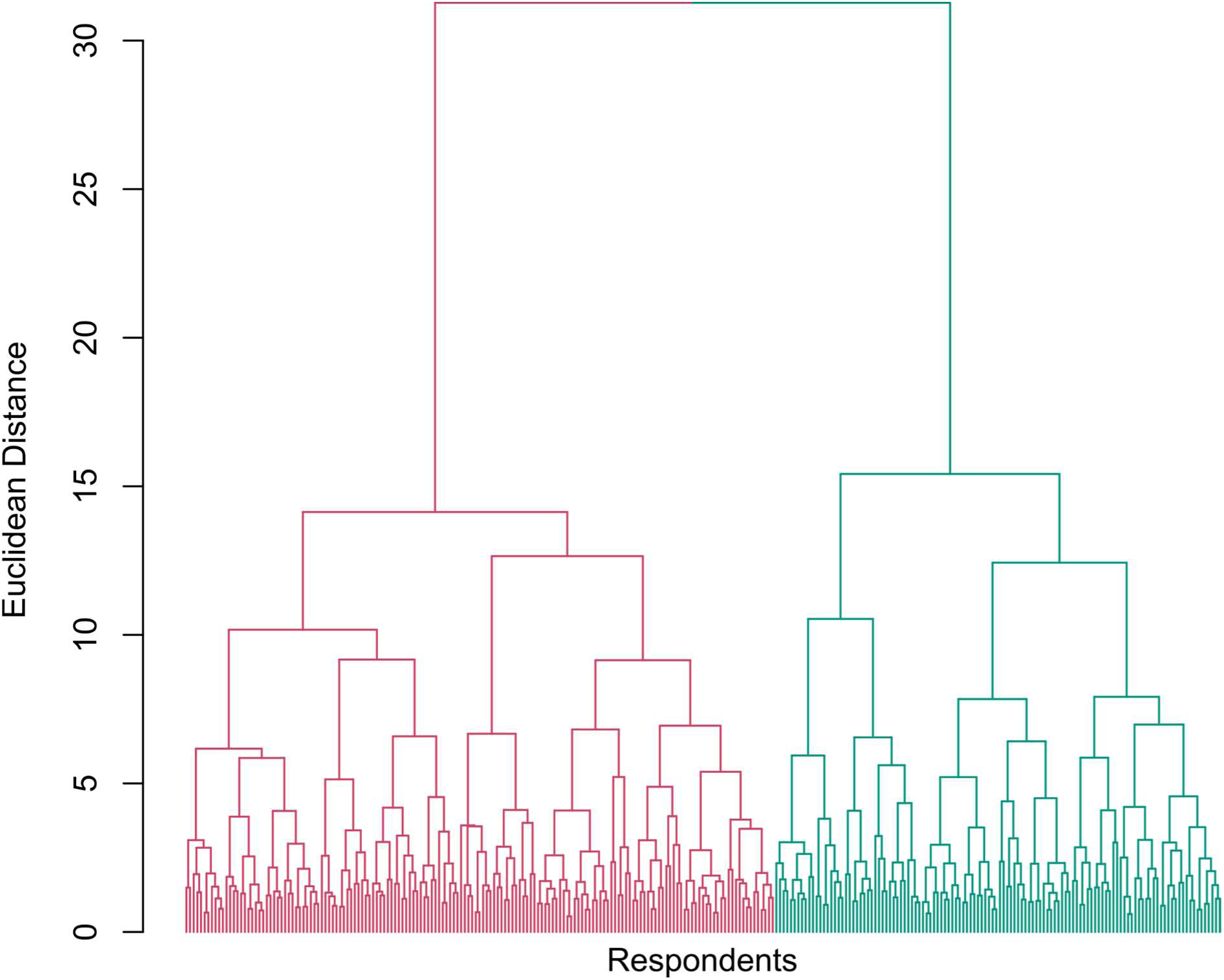
Dendogram of the cluster analysis. The colouring (pink, green) illustrates the two- cluster solution.

**Figure 20.**
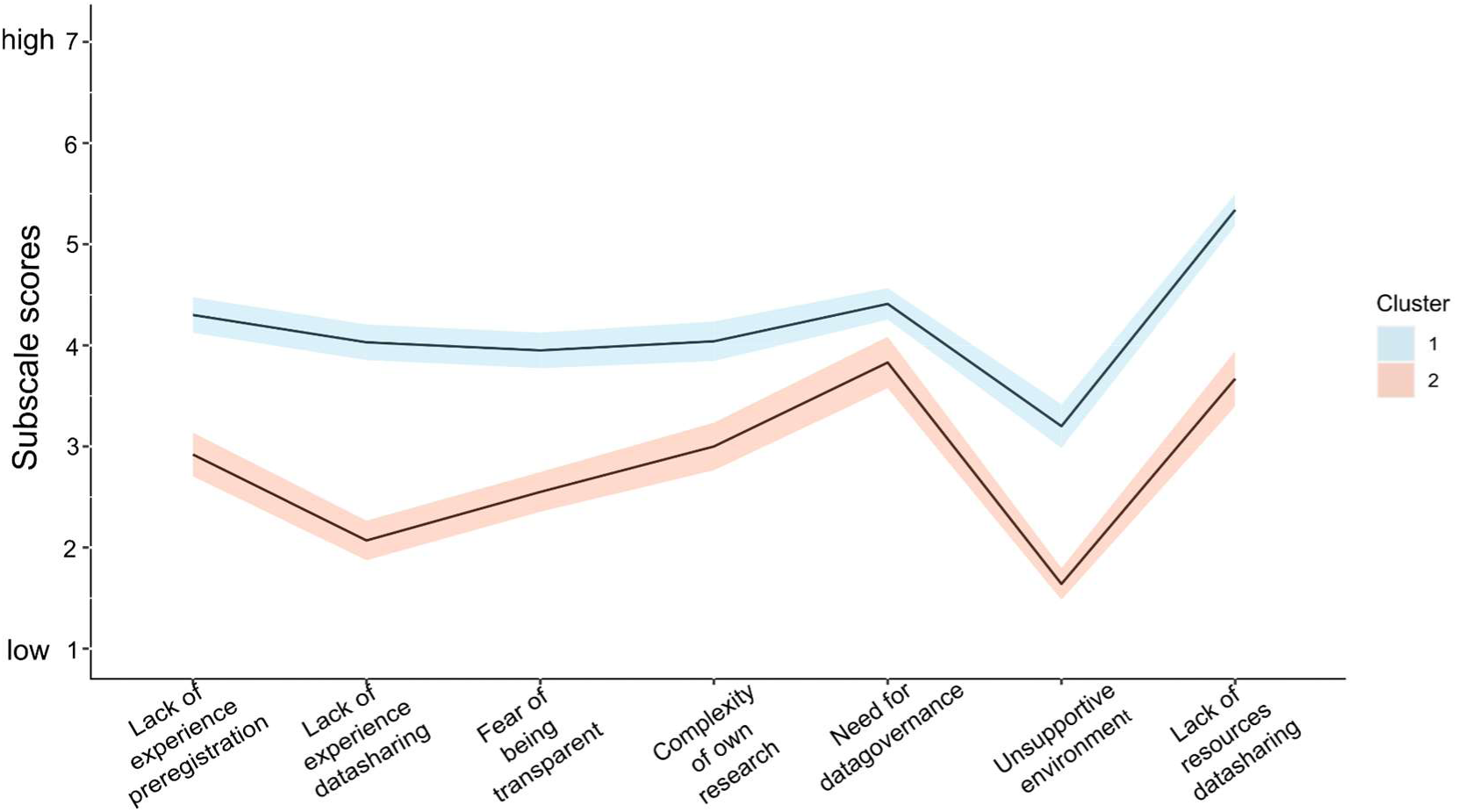
Profile plot of subscale scores for clusters 1 and 2. Coloured area shows 95% confidence intervals.

To find out whether cluster-belongingness could be explained by demographic variables, we conducted a regression analysis. Overall the explanatory power of our regression model was marginally better than chance, *Χ*²(4)=10.09, p=0.039, (Table 4) with an out-of-sample accuracy of 59,9%, based on 10-fold cross-validation. The affiliation with a medical faculty and full/associate professorship predicted whether a participant belonged to cluster 1 at trend level, with *p*=0.059 and *p*=0.067, respectively.

**Table 4.** . Results from logistic regression with Cluster as the dependent variable and the demographic variables “research experience”, “career level”, “EU residency” and “Affiliation with medical faculty” as predictors.

| Variable | Estimate | Std. Error | z-value | p-value |
| --- | --- | --- | --- | --- |
| (Intercept) | -0.329 | 0.330 | -0.996 | 0.319 |
| Research experience | 0.113 | 0.312 | 0.361 | 0.718 |
| Career level | -0.570 | 0.312 | -1.830 | 0.067 |
| EU residency | 0.082 | 0.267 | 0.306 | 0.759 |
| Affiliation with medical faculty | 0.498 | 0.264 | 1.884 | 0.059 |

## 4 Discussion

Preregistration of research questions, hypotheses and the analysis plan as well as data sharing were proposed to improve the replicability, robustness and reproducibility ^16, 39^. This survey aimed to shed light on the experience with and attitude towards open science practices in human neuroimaging, namely with regards to preregistration, data sharing and data standards. We reached out to researchers who had published papers using human fMRI in the past, which was reflected by the resulting sample being mainly composed of researchers who were advanced in their careers. It can be assumed that most participants of this survey were heading their own labs and that they oversaw and exerted influence in their field of research.

Surprisingly, the interest to use preregistration was rather modest. About one half of participants had preregistered a study before, with OSF as the most commonly used platform. There was no indication of a trend towards more widespread use of preregistration in the future. Still, two thirds had at least thought about preregistering their research. Besides the barriers and fears that we had asked for, some participants shared a critical perspective on the role of preregistration as a technique to promote the quality of science (Table 5, cf. ^17, 40^). This view stands against advocates of preregistration who see no alternative to prevent hindsight bias and overconfidence in research findings ^14, 41^. Best practice guidelines explaining when and how to preregister neuroimaging research, which is often exploratory and complex, have not been established, yet, although new templates such as PRP-QUANT ^42^ and an OSF- template were made available, which is an important step in this direction. Furthermore, this survey demonstrated the rising awareness for the importance of data sharing in the neuroimaging community. Most participants had thought about online data sharing before. Data sharing mechanisms with access governance were clearly prefered (c.f. ^43^ ), while one third of participants also preferred sharing with unrestricted access. At large, the results are in line with the findings from Houtkoop et al.^7^ who surveyed psychologists about their views on data sharing practices. Although comparisons between their results and ours remain descriptive and are somewhat limited because of differences in methods, we observed some differences regarding data protection concerns: Compared to psychologists, neuroimaging researchers more frequently reported that their institutional review boards prohibited data sharing (30% vs. 5%), that they were constrained by lacking explicit consent from subjects to share data (41% vs. 28%), and that anonymity cannot be guaranteed (45% vs. 16%). Lastly, While one third of respondents were using BIDS in current neuroimaging projects, we observed a strong interest to adopt BIDS in the future. Nonetheless, a substantial proportion of participants had not yet heard about BIDS. The major bottleneck for adopting BIDS appears to be limited time. This finding may reflect the expectation that introducing a new data standard to the lab would cost a lot of resources. Such apprehension is understandable in the face of limited resources that are available for research in the public domain. The availability of software that is easier to operate, e.g. to convert data into BIDS via GUI instead of command line interface, may facilitate the implementation of the data standard in more labs with less experience in programming (c.f. Table 5).

Fears and barriers in the way of adopting open science practices may be governed by a few underlying dimensions. If the latter were known, further research could investigate how these factors are shaped by current research practices, whether they relate to certain parameters (e.g. demographic variables), and whether they are amenable to targeted intervention. In a first approach to this question, we identified seven factors driving the responses to this survey. An exploratory analysis of these factors revealed some interesting differences between sub-groups of participants: experienced researchers at lower career level expressed higher fears of being transparent as compared to those at higher career level. It may be speculated that post-doctoral researchers are particularly anxious to be outpaced by their peers and therefore cultivate a less transparent and more protective style to maximize their chances in the race for a small number of tenured positions available in academia. Once they have achieved a tenured position, professors may be more relaxed to let other researchers work with their data and may have less fear that preregistration may make them vulnerable to lose in the competition for resources. In the race for tenured positions, post-docs may also be inclined to drop the extra work to meet open science standards, because many academic employers and funders still weigh open science practices less as compared to traditional measures of scientific output such as impact factors and the number of publications (c.f. Table 5). Furthermore, a higher need for data governance was expressed by researchers at medical faculties as well as researchers residing within the EU, aligning with higher agreement among EU residents that they were not allowed to share imaging data and lower agreement to share primary data from their next neuroimaging study online. Research with patients in general is subject to strict juridical regulations for data protection. The GDPR has increased data protection requirements recently and caused irritation among researchers across Europe about how to reconcile data protection regulations with the sharing of human data. However, comparisons of EU vs. non-EU participants should be taken with some caution, as we did not actively match the groups based on demographic characteristics. Researchers at medical faculties as compared to other faculties also had higher chances to face an unsupportive environment in terms of adopting reproducibility practices. The dual workload of clinical work and research, often paired with the pressure to produce high-ranking research output, is not suited to create an environment where scientist practitioners engage in new techniques. To score high on the factor “unsupportive environment” one had to indicate that one’s supervisor would not support reproducibility practices. Professors naturally scored low on these items.

We aimed to test the existence of distinct subgroups which differed in their profiles on open science fears and barriers. If such subgroups were identified, and generalized to a larger research community, this could inform how more targeted interventions, teaching programs, and policies can be developed. Cluster analysis revealed two groups that were either characterized by generally higher or generally lower fears and barriers. This suggests that while a large portion of the neuroimaging community is well-versed in open science practices, an equally large portion is lagging behind and might benefit from broad awareness and teaching programs, comprising all aspects of open science. To take the concerns of this group seriously, the community should work out detailed guidelines to reconcile preregistration with challenges brought along by complex imaging projects and dominantly explorative research. Further instruments to respond to the many barriers and fears of data sharing have been described elsewhere ^7^. Explorative regression analysis showed that the demographic variables we had used to predict belongingness to the two clusters barely exceeded chance level and the out-of-sample accuracy was relatively low. None of the variables that were tested predicted cluster belongingness beyond trend level. Future research is necessary to confirm our findings and to explore more variables that may aid the prediction.

### 4.1 Limitations

Conclusions from this survey are limited by the low response rate to the survey invitation (2.4%), which was below the rates reported in previous investigations (4%^27^, 5%^7^, 9%^24^). Studies like ours that remove incomplete responses tend to find lower response rates. In addition, unlike previous studies, we did not recruit via our professional networks. While the latter is an effective strategy to increase responses, it may have the drawback of inflating the proportion of participants sharing a certain perspective on the topic (although the recruitment strategy we used does not protect against that bias). Also, the pandemic situation and the increase in unsolicited survey invitations we are observing in recent years may have had a negative impact on the willingness to participate in our study. Clearly, researchers would only take the effort to participate when they shared a basic interest in the topic. The cluster analysis showed that about half of the sample reported less experience with and training in open science techniques as well as higher fears thereof, and we found that about 41% did not know BIDS before. Although the sample may not be representative, these numbers evidence reasonable variance in familiarity and attitude towards open science practices, which is necessary to receive meaningful results. The recruitment strategy emphasized on researchers working with fMRI and investigating humans, generalization to researchers working with other neuroimaging modalities and other species is therefore limited. BIDS was initially introduced for human fMRI, therefore the results from this sample are easier to interpret as compared to a more heterogeneous sample of researchers working with different modalities, for whom the data standard became available later or which were not yet covered by BIDS at the time of this survey taking place. As we approached researchers who had published as corresponding authors before, conclusions cannot be generalized to very early career researchers. Also, it should be noted that the questionnaire we had used is not a validated instrument. The Open Scholarship Survey^44^, for instance, which has been designed for the investigation of similar research questions as ours, was not yet available when this project was started. Thus, the factor analytic results need to be interpreted in the context of this survey. We focused on barriers and fears, and did not interrogate beliefs about the benefits of open science (e.g. that open science practices can increase the quality and impact of one’s research output). Also, we did not assess objective measures such as the number of preregistered studies or the number of shared data sets, information that could be used for validation. Finally, some aspects of open science were not touched by the survey such as sharing of materials and code. Thus, the results cover certain aspects of open science practices while others are not illuminated. Finally, a few ambiguities in the questions where discovered by the participants who had shared their feedback with us (c.f. Table 5).

### 4.2 Conclusions

Limited time and insufficient education about tools to structure and share data were reported as the major barriers for adopting open science practices. Although half of the participants were experienced with preregistration, the willingness to preregister studies in the future was restrained, and some participants expressed a rather critical view on preregistration. Neuroimaging researchers are open to data sharing and most have experience with sharing primary research data. Concerns regarding the protection of the privacy of participants from neuroimaging experiments and missing sections in consent forms to enable data sharing (cf. ^45^) make researchers hesitant to share neuroimaging data. Measures to reinforce data sharing, to educate researchers how to prepare consent forms enabling data sharing, and to inform about existing infrastructure and mechanisms of data protection may increase the willingness to share primary neuroimaging data. Analyses of individual differences suggest that some groups of researchers may benefit more from certain measures to facilitate the usage of open science techniques: (1) Experienced researchers before tenure may benefit from measures reducing fears of being transparent. (2) Researchers in the EU may benefit from measures to satisfy the need for data governance. (3) Researchers at medical faculties would also benefit from measures to satisfy the need for data governance. In addition, (4) they would benefit from measures aiming to create an environment that is more supportive of open science practices.

## 4.3 CRediT authorship contribution statement

CP: Conceptualization, Methodology, Verification, Data Curation, Supervision, Project administration, Writing - Original Draft.

NU, MS: Software, Formal analysis, Data Curation, Writing - review & editing. FF, RP, CS: Methodology, Writing - review & editing.

MS: Investigation, Project administration.

## 4.5 Acknowledgement

Thanks are due to Gordon Feld for critical reading of an earlier version of this manuscript. We are thankful to our colleagues from the Department for Psychosomatic Medicine and Psychotherapy, CIMH, for their feedback on the questionnaire during development.

## Comments on further barriers in the way of open science

*General*

- Not forwarding career of aspiring PI
- Engineer would be needed for implementation

**BIDS**

- Some format aspects such as tsv make BIDS inconvenient to use
- Journals require posting of primary data in idiosyncratic format, not in BIDS
- No MATLAB based option to convert to BIDS available

**Preregistration**

- Difficulties getting preregistered report on longitudinal data accepted because first wave already collected
- Pre-registered analyses are often outdated once the study is complete
- Research questions that we address are always against the limits of what current analysis tools are capable of doing; questions mostly requires fine-tuning methods, developing new approaches, bringing in other tools, etc.
- Preregistration constrains the creativity that is at the basis of progress in science
- Preregistration leads to terrible papers, where too much text is spent on explaining the preregistered content and the justifications for deviating from them
- Realistic standards for evaluating conformity to the preregistration missing
- Pre-registration is only meaningful for purely confirmatory studies. Purely confirmatory studies are only meaningful when there is a strong hypothesis and the goal of the confirmatory study is to confirm this hypothesis.
- The benefits of pre-registration have not been thoroughly demonstrated in order to merit its adoption

**Data sharing**

- Data protection regulations from host institution incompatible with sharing
- Money to store and manage data repositories missing after grant terminates
- Neuroimaging data are intellectual property, rights of researchers acquiring data need to be protected
- No canonical interpretation of the laws/regulations available
- Practical guides on how to share clinical data online missing
- Whether the data will be used by anyone at all, and how long a given repository will last is unknown.

## Comments expressing further fears of open science

- Lose my job because not complying with host institutions data protection regulation
- My worries about not being able to publish every last ounce of results from my data are very high.
- I unfortunately think that the open science movement has the capacity to really disadvantage jr. researchers in comparison to well-established labs
- Transparency is nice, but we seem to be willing to sacrifice part of our creativity through forced standardization
- My greatest fear is giving away your research ideas with preregistration

## Feedback on the questionnaire

- Don’t think this survey captured my opinions very accurately. I am a strong supporter of Open Science, but have a number of concerns about data sharing and the potential for abuse
- A question was lacking about lack of confidence in how to interpret the jurdical bases for data sharing
- In the survey it was a bit unclear if data sharing refers to neurogimaging data only or in general
- Many researchers will not reply, let alone reply honestly
- I think that analyses for individual papers can be prespecified, but it would be hard to pre-specify analyses for large studies. I understood that you are referring to pre-registration of the entire large study, which I said I do not do
- There was insufficient opportunities to comment on the role of journals (static, laminated publications etc) in effectively prohibiting open science practices. Open science may obviate the need for journals.
- The question at the bottom of the page asking for legal issues yes/no was difficult to answer, because we have these issues for old data (not considering data sharing) but we always take care of these now in new projects (including data sharing).
- Many of your questions are difficult to answer / ambiguous since there are different hurdles to share data from healthy participants and patients

## Other

Preregistration provides a way of claiming precedence for an idea, even if the results don’t bear out the findings

Table 5. In the end of the survey, the participants were given the opportunity to write a free-text comment to the authors of the survey. 45 (17%) of the participants took advantage of this option. The table lists a selection of these comments that bring up aspects that were not properly covered by the survey questions, or that give constructive feedback on the questionnaire itself. Comments have been shortened or reworded at the discretion of the author (CP) to make them more concise..

## References

1. Botvinik-Nezer, R. et al. Variability in the analysis of a single neuroimaging dataset by many teams. Nature 1–7 (2020) doi:10.1038/s41586-020-2314-9.

2. Poldrack, R. A. et al. Scanning the horizon: towards transparent and reproducible neuroimaging research. Nat. Rev. Neurosci. 18, 115–126 (2017).

3. Eklund, A., Nichols, T. E. & Knutsson, H. Cluster failure: Why fMRI inferences for spatial extent have inflated false-positive rates. Proc. Natl. Acad. Sci. U.S.A. 113, 7900– 7905 (2016).

4. Button, K. S. et al. Power failure: why small sample size undermines the reliability of neuroscience. Nat Rev Neurosci 14, 365–376 (2013).

5. Guo, Q. et al. The reporting of observational clinical functional magnetic resonance imaging studies: a systematic review. PLoS ONE 9, e94412 (2014).

6. Carp, J. The secret lives of experiments: Methods reporting in the fMRI literature. NeuroImage 63, 289–300 (2012).

7. Houtkoop, B. L. et al. Data Sharing in Psychology: A Survey on Barriers and Preconditions. Advances in Methods and Practices in Psychological Science 1, 70–85 (2018).

8. Valizadeh, S. A., Liem, F., Mérillat, S., Hänggi, J. & Jäncke, L. Identification of individual subjects on the basis of their brain anatomical features. Sci Rep 8, 5611 (2018).

9. Wachinger, C. et al. BrainPrint: a discriminative characterization of brain morphology. Neuroimage 109, 232–248 (2015).

10. Amico, E. & Goñi, J. The quest for identifiability in human functional connectomes. Sci Rep 8, 8254 (2018).

11. Bari, S., Amico, E., Vike, N., Talavage, T. M. & Goñi, J. Uncovering multi-site identifiability based on resting-state functional connectomes. Neuroimage 202, 115967 (2019).

12. Finn, E. S. et al. Functional connectome fingerprinting: identifying individuals using patterns of brain connectivity. Nat Neurosci 18, 1664–1671 (2015).

13. Gorgolewski, K. J. et al. The brain imaging data structure, a format for organizing and describing outputs of neuroimaging experiments. Sci Data 3, 160044 (2016).

14. Nosek, B. A., Ebersole, C. R., DeHaven, A. C. & Mellor, D. T. The preregistration revolution. Proc Natl Acad Sci USA 115, 2600 (2018).

15. Ioannidis, J. P. A. Why most published research findings are false. PLoS Med. 2, e124 (2005).

16. Munafò, M. R. et al. A manifesto for reproducible science. Nature Human Behaviour 1, 1–9 (2017).

17. Gelman, A. & Loken, E. The Statistical Crisis in Science. American Scientist 102, 460– 465 (2014).

18. Carp, J. On the plurality of (methodological) worlds: estimating the analytic flexibility of FMRI experiments. Front Neurosci 6, 149 (2012).

19. Gentili, C., Cecchetti, L., Handjaras, G., Lettieri, G. & Cristea, I. A. The case for preregistering all region of interest (ROI) analyses in neuroimaging research. European Journal of Neuroscience 53, 357–361 (2021).

20. Esteban, O. et al. fMRIPrep: a robust preprocessing pipeline for functional MRI. Nat. Methods 16, 111–116 (2019).

21. Nichols, T. E. et al. Best practices in data analysis and sharing in neuroimaging using MRI. Nat Neurosci 20, 299–303 (2017).

22. Gorgolewski, K. J. et al. BIDS apps: Improving ease of use, accessibility, and reproducibility of neuroimaging data analysis methods. PLOS Computational Biology 13, e1005209 (2017).

23. Laird, A. R. Large, open datasets for human connectomics research: Considerations for reproducible and responsible data use. NeuroImage 244, 118579 (2021).

24. Tenopir, C. et al. Data Sharing by Scientists: Practices and Perceptions. PLOS ONE 6, e21101 (2011).

25. White, T., Blok, E. & Calhoun, V. D. Data sharing and privacy issues in neuroimaging research: Opportunities, obstacles, challenges, and monsters under the bed. Hum Brain Mapp (2020) doi:10.1002/hbm.25120.

26. Sayogo, D. S. & Pardo, T. A. Exploring the determinants of scientific data sharing: Understanding the motivation to publish research data. Government Information Quarterly 30, S19–S31 (2013).

27. Schmidt, B., Gemeinholzer, B. & Treloar, A. Open Data in Global Environmental Research: The Belmont Forum’s Open Data Survey. PLOS ONE 11, e0146695 (2016).

28. Poline, J.-B. et al. Data sharing in neuroimaging research. Front Neuroinform 6, 9 (2012).

29. Paret, C. Survey on Open Science-Practices in Functional Neuroimaging. Dataset and Materials. https://github.com/christianparet/Survey-on-Open-Science-Practices-in-Functional-Neuroimaging.-Dataset-and-Materials (2021) doi:10.5281/zenodo.5729056.

30. Leiner, D. J. SoSci Survey. (2019).

31. Morey, R. D., et al. BayesFactor: Computation of Bayes Factors for Common Designs. (2018).

32. Rosseel, Y. lavaan: An R Package for Structural Equation Modeling. Journal of Statistical Software 48, 1–36 (2012).

33. Marsh, H. W., Morin, A. J. S., Parker, P. D. & Kaur, G. Exploratory Structural Equation Modeling: An Integration of the Best Features of Exploratory and Confirmatory Factor Analysis. Annual Review of Clinical Psychology 10, 85–110 (2014).

34. Kassambara, A. & Mundt, F. factoextra: Extract and Visualize the Results of Multivariate Data Analyses. (2020).

35. Kuhn, M. Building Predictive Models in R Using the caret Package. Journal of Statistical Software 28, 1–26 (2008).

36. Chambers, C. D., Dienes, Z., McIntosh, R. D., Rotshtein, P. & Willmes, K. Registered reports: realigning incentives in scientific publishing. Cortex 66, A1–2 (2015).

37. Esteban, O. et al. MRIQC: Advancing the automatic prediction of image quality in MRI from unseen sites. PLOS ONE 12, e0184661 (2017).

38. Markiewicz, C. J. et al. OpenNeuro: An open resource for sharing of neuroimaging data. 2021.06.28.450168 https://www.biorxiv.org/content/10.1101/2021.06.28.450168v2 (2021) doi:10.1101/2021.06.28.450168.

39. Nosek, B. A., et al. Replicability, Robustness, and Reproducibility in Psychological Science. (2021).

40. Szollosi, A. et al. Is Preregistration Worthwhile? Trends in Cognitive Sciences 24, 94–95 (2020).

41. Nosek, B. A., et al. Preregistration Is Hard, And Worthwhile. Trends Cogn Sci 23, 815– 818 (2019).

42. Preregistration Task Force. Preregistration Standards for Psychology - the Psychological Research Preregistration-Quantitative (aka PRP-QUANT) Template. (2021) doi:10.23668/PSYCHARCHIVES.4584.

43. Cheah, P. Y., et al. Perceived Benefits, Harms, and Views About How to Share Data Responsibly: A Qualitative Study of Experiences With and Attitudes Toward Data Sharing Among Research Staff and Community Representatives in Thailand. Journal of Empirical Research on Human Research Ethics 10, 278–289 (2015).

44. Center for Open Science. Open Scholarship Survey. https://www.cos.io/initiatives/open-scholarship-survey.

45. Bannier, E. et al. The Open Brain Consent: Informing research participants and obtaining consent to share brain imaging data. Human Brain Mapping 42, 1945–1951 (2021).

